# Mechanisms of ion selectivity and rotor coupling in the bacterial flagellar sodium-driven stator unit

**DOI:** 10.1101/2022.11.25.517900

**Authors:** Haidai Hu, Philipp F. Popp, Mònica Santiveri, Aritz Roa-Eguiara, Yumeng Yan, Zheyi Liu, Navish Wadhwa, Yong Wang, Marc Erhardt, Nicholas M. I. Taylor

## Abstract

Bacteria swim using a flagellar motor that is powered by stator units. These stator units are energized by an ionic gradient across the membrane, typically proton or sodium. The presumed monodirectional rotation of the stator units allows the bidirectional rotation of the flagellar motor. However, how ion selectivity is attained, how ion transport triggers the directional rotation of the stator unit, and how the stator unit is incorporated into the motor remain largely unclear. Here we have determined by cryo-electron microscopy the structure of the Na^+^-driven type stator unit PomAB from the gram-negative bacterium *Vibrio alginolyticus* in both lipidic and detergent environments, at a resolution up to 2.5 Å. The structure is in a plugged, auto-inhibited state consisting of five PomA subunits surrounding two PomB subunits. The electrostatic potential map uncovers sodium ion binding sites within the transmembrane domain, which together with functional experiments and explicit solvent molecular dynamics simulations, suggest a mechanism for ion translocation and selectivity. Resolved conformational isomers of bulky hydrophobic residues from PomA, in the vicinity of key determinant residues for sodium ion coupling of PomB, prime PomA for clockwise rotation. The rotation is tightly blocked by the trans-mode organization of the PomB plug motifs. The structure also reveals a conformationally dynamic helical motif at the C-terminus of PomA, which we propose regulates the distance between PomA subunit cytoplasmic domains and is involved in stator unit-rotor interaction, concomitant stator unit activation, and torque transmission. Together, our studies provide mechanistic insight for understanding flagellar stator unit ion selectivity and incorporation of the stator units into the motor.

## Introduction

Many bacteria rotate flagella to power their movement. The flagellum is characterized by a long filament, connected through a flexible hook to cell envelope embedded rotary motor (or basal body), which comprises a rotor and multiple stator units^1–4^. The flagellar stator unit uses the transmembrane ion motive force (IMF) to generate mechanical torque to rotate the flagellum, which is employed by many bacteria to direct their locomotion in liquid environment or on viscous surfaces to a favorable niche^4–6^. Driven by the stator unit, the bacterial flagellar motor can rotate in both clockwise (CW) and counterclockwise (CCW) directions, with the switch between the two directions controlled by intracellular chemotaxis signaling ^7,8^. The stator units are strictly required for rotation of the flagellum and thus motility of the bacteria, but not for flagellar assembly^9,10^. In addition, the stator units dynamically associate with and dissociate from the rotor^11–13^. Changing the number of engaged stator units allows tuning the required torque in relation to the mechanical load ^14–18^.

Each stator unit is composed of two membrane proteins assembled as a complex buried inside the cytoplasmic membrane, in which their transmembrane domains organize as an ion channel^19,20^. Incorporation of the stator unit requires its cytoplasmic domain to interact with the rotor and its periplasmic domain to attach to the bacterial cell wall^21^. During activation, the stator unit undergoes a conformational change from a plugged state into an unplugged one, and the subsequent ion translocation through the stator unit drives its activity^22,23^. Hence, the stator unit itself is considered a “miniature” motor. Torque generated by ion translocation is transmitted to the rotor via electrostatic interactions at the stator-rotor interface^24–27^. Depending on the conducting ions, stator units can be grouped into two subfamilies: H^+^-driven stator unit (e.g., MotAB) and Na^+^-driven stator unit (e.g., PomAB)^28,29^. In addition, stator units use potassium and divalent ions such as calcium or magnesium as coupling ions have also been reported^30–33^. However, at the molecular level, how stator units discriminate among different types of ions and power rotation of the flagellar motor have remained unclear.

Recently, single particle cryo-electron microscopic (cryo-EM) structures of H^+^-driven MotAB stator units^34,35^, cryo-EM structures of intact flagellar motor complexes^36–38^, as well as *in situ* cryo-electron tomographic (cryo-ET) studies of the flagellar motor^21,39–41^, provided detailed structural and functional views of stator unit assemble, torque generation and motor function ^1^. The data strongly suggest a rotational model for the mechanism of action of the stator units. Upon dispersion of the IMF, MotA is proposed to rotate around MotB, which is anchored to the peptidoglycan layer. By engaging with the rotor MotA rotation powers the rotation of the large rotor. The differential engagement of MotA with the rotor between the CW and CCW states of the rotor is proposed to form the mechanistic basis of switching.

Cryo-EM reconstructions of the Na^+^-driven stator unit PomAB from *V. alginolyticus* (*Va*PomAB) and *V. mimicus* have also been reported^34,35^. However, due to the low resolution and anisotropic maps, the atomic coordinates of the Na^+^-driven stator unit remain unknown. The Na^+^-driven stator unit is particularly important for *Vibrio* species, including pathogenetic ones, as their polar flagella can only be powered by the transmembrane Na^+^ gradient. Furthermore, the Na^+^-driven stator unit is an ideal subject for investigating stator unit ion selectivity and translocation mechanisms. As a Na^+^ ion interacts more with electrons than a proton in the cryo-electron microscope, it could potentially be visualized more readily in a high-resolution cryo-EM map. Finally, sodium ions are easier to be detected and manipulated than protons^42^.

The atomic structure of the Na^+^-driven stator unit is thus crucial for the mechanistic understanding of how the stator unit distinguishes ions and couples ion transportation into its rotation. To this end, we determined cryo-EM structures of *Va*PomAB in both detergent and lipidic environments, with the local map resolution reaching up to ∼2 Å. The high-resolution structure enabled us to locate Na^+^ ion binding sites and revealed the structural and mechanistic basis of the ion selectivity. We show at the molecular level how the stator unit achieves its monodirectional rotation upon ion transport. Furthermore, we identified a helical motif C-terminal of PomA (CH) that is crucial for stator unit function. We validated our structural results through extensive mutagenetic analysis and molecular dynamics (MD) simulations. Finally, we propose a role for the asymmetric cytoplasmic domain arrangement of the stator unit in the torque generation and the assembly and disassembly mechanism of the stator unit into the motor.

## Results

### Structure determination and overall architecture of *Va*PomAB

Intact *Va*PomAB is an anisotropically shaped complex and shows preferential orientation of particles in vitreous ice^35^. To improve sample homogeneity, we modified the protein purification protocol and encoded a protease site in the PomB gene after the plug region, which allowed for the removal of the PomB peptidoglycan binding domain (PGB) during protein purification. To overcome the preferred orientation, we added the zwitterionic detergent CHAPSO to randomize particle orientation during EM grid preparation^43,44^. Single-particle analysis yielded an overall resolution of *Va*PomAB in LMNG detergent of approximately 2.5 Å resolution, with the cryo-EM map of sufficient quality to build an atomic model for most of the protein complex. The local resolution corresponding to the inner transmembrane domain approaches to 2 Å, with clear density for non-protein molecules, allowing us to model water molecules and ions, as well as residue side chains isomers (Fig. 1, Extended Data Fig. S2-S4 and Table S1).

**Fig. 1.**
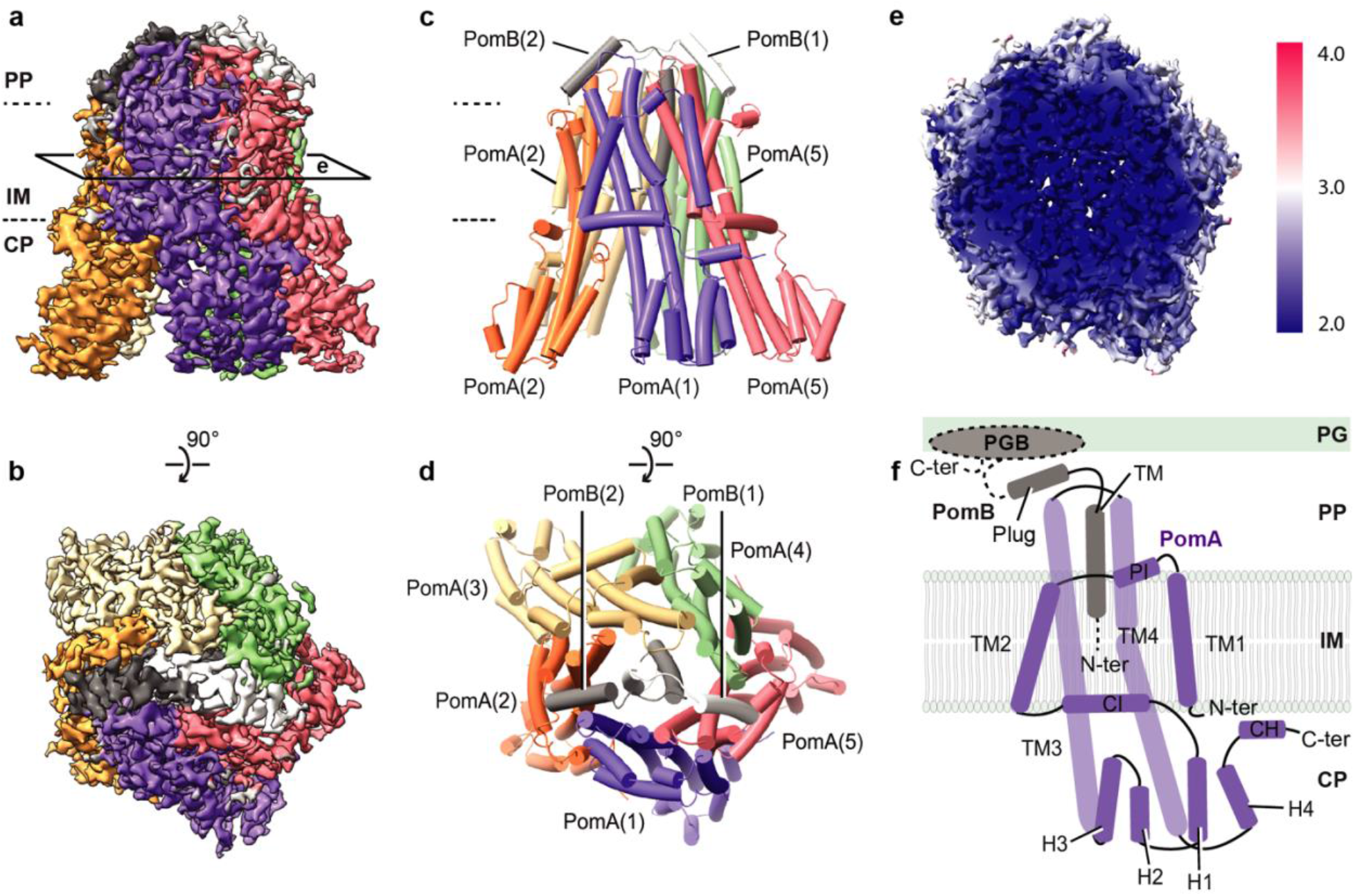
Cryo-EM map and overall architecture of the Na^+^-driven stator unit *Va*PomAB. **a**, Cryo-EM map of *Va*PomAB. PomA subunits (purple, orange, yellow, green and red) surround PomB subunits (black and white) viewed from the plane of the membrane. Dashed lines represent approximate inner membrane boundaries. **b**, Cryo-EM map of *Va*PomAB viewed from the periplasmic side. **c**, Ribbon model representation of *Va*PomAB. Subunits are colored as in **a. d**, *Va*PomAB model viewed from the periplasmic side. **e**, Local resolution map of *Va*PomAB viewed from a cross section as indicated in **a. f**, Topology diagram and secondary structural elements of *Va*PomA (purple) and *Va*PomB (black) subunits. The gray ellipse indicates the PomB peptidoglycan-binding domain (PGB). Abbreviations: PP, periplasm; IM, inner membrane; CP, cytoplasm; PG, peptidoglycan; TM, transmembrane; H, helix.

PomAB assembles the characteristic bell shape of the stator unit family, with conserved 5:2 subunit stoichiometry and overall architecture. Five PomA molecules arrange pseudo-symmetrically around two PomB, with each PomA subunit comprising four transmembrane helices (TM1-TM4) folded into two radial layers. The TM3s and TM4s form an inner layer lining the dimerized PomB TMs. The TM1s and TM2s surround peripherally, together with PomA periplasmic interface (PI) helices and cytoplasmic interface (CI) helices, establishing an outer layer, with one side packing against the inner layer and the other side hydrophobically interacting with the lipid bilayer. The resolved TM1 of one PomA makes prominent contact with the TM2 from the adjacent subunit. The cytoplasmic domain of PomA contains a compact helix bundle (H1-H4), a region where torque is generated through electrostatic matching with the rotor FliG torque helix^45^. The cryo-EM map of PomAB also reveals a short helix after PomA H4, which we designated as CH (C-terminal helix) motif, attaching to the CI helix of a neighboring PomA subunit (Fig. 1f). The plugged motifs from two PomB chains are fully resolved in our PomAB structure, where they are positioned on the top of the periplasmic side of the stator unit, consistent with a plugged autoinhibited state. We also noticed that each end of the plug motif interacts with the PI helix of PomA. We propose this causes the N-terminal residues (residues 1–21) of two PomA subunits to be disordered, as these are not resolved in our cryo-EM map (Extended Data Fig. S5c).

### Plug motif and autoinhibition mechanism

The PomB plug motif is a short amphipathic α-helix, following the TM helix. Deletion of the plug motif results in ion influx into the cell cytosol, causing cell growth inhibition when overexpressed^23,46^. Earlier studies through mutagenesis and cross-linking experiments have mapped critical residues involved in interactions between the PomB plug motif and PomA^23,47^. In the PomAB structure, the TM of PomB is connected to the plug helix through a four-residue linker (Fig. 2c), which makes the plug helix turn approximately 145°, rendering its C-terminus to point towards the cytoplasmic membrane. The two short linkers establish interaction laterally by four backbone hydrogen bonds, organizing the plug motifs as a trans-mode configuration relative to PomB TM helices, with a pseudo-mirror symmetry perpendicular to the cell membrane (Fig. 2a, 2d). The plug motifs from sodium and proton-driven stator units share a similar amino acid pattern (Extended Data Fig. S1b) comprising a hydrophobic side that makes its main interaction with the stator unit itself and a hydrophilic side that is exposed in most parts to the periplasmic space solvent (Fig. 2b). The contact environment of the PomB plug motif is mainly contributed by a cleft framed by the periplasmic side of the TM4, TM3 and the PI helix from one PomA subunit, and the periplasmic side of the TM3 and TM4 from the adjacent PomA subunit. Three residues from the plug motif (I50, M54 and F58) deeply insert into this cleft, establishing hydrophobic interactions (Fig. 2e-f). Additionally, the PomB F47 aromatic ring is sandwiched between the pyrrolidine ring of P172 and the side chain of M169 from PomA, via CH-π interactions, further stabilizing the plug motif (Fig. 2f). The 5:2 stoichiometry of the stator unit creates inequivalent binding environments for the two plug motifs, as examined by calculating the surface buried area and free energies of residues forming the plug helix (residues 44-58, Fig. 2g-h). Therefore, we speculate that during stator unit activation, releasing the plug motif from the stator unit is not a symmetric process. Instead, one plug motif with relatively low binding energy likely detaches from its inhibitory site first, and the second plug motif will then be induced to be released. PomB G59 marks the end of the plug motif, and it directly exerts the effect on the conformation of PomA PI helix. We found that each of the PomB plug motifs induces two different conformations of the PomA PI helix that links PomA TM1 and TM2; one conformation is akin to those observed in the other three PomA subunits, and the other conformation extends TM2 one more helical turn involving residues from L26 to V32 (Extended Data Fig. S5c).

**Fig. 2.**
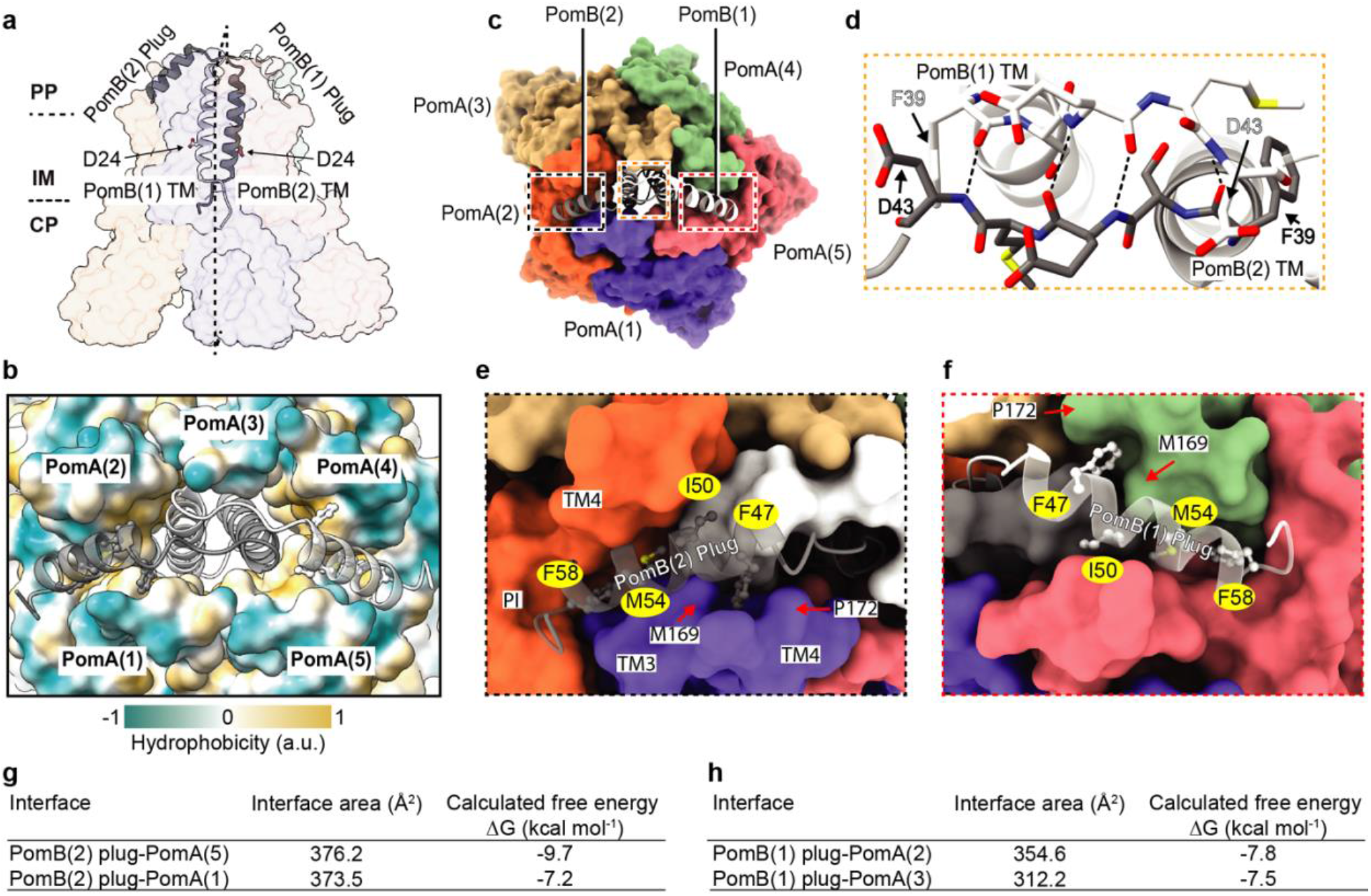
PomB plug motif and auto-inhibition mechanism. **a**, *Va*PomAB in its auto-inhibited state, viewed from the plane of the membrane, with PomB shown as ribbons (black and white) and PomA shown as a semitransparent surface representation. The aspartate residues D24 from both PomB TM are indicated and shown as sticks. **b**, Top view of *Va*PomAB with PomB shown as ribbons and PomA shown as a surface representation colored according to its hydrophobicity. **c**, Top view of *Va*PomAB. PomA subunits are shown as a surface representation and PomB subunits are displayed as ribbons, colored as in Fig. **1a. d**, Close-up view from the periplasmic side of the interactions of the linkers (Phe39-Asp43) that connect PomB plug motifs and TMs (it corresponds to the yellow box in **c**). Hydrogen bonds are represented as dashed lines. **e**, Plug motif from PomB(2) binding environment (black box in **c**). **f**, Plug motif from PomB(1) binding environment (red box in **c**). **g**-**h**, Calculated interface buried area and free energy of PomB plug motifs.

The dynamics of the PomA PI helices stemming from the PomB plug motif interaction presumably drives the flexibility of the corresponding TM1, as the latter could not be resolved in two of the PomA subunits. The high-resolution PomAB structure was determined in a detergent micelle environment, raising the possibility that detergent molecules could have an impact on the conformation of PomAB, particularly the membrane-facing helices, including the disordered TM1 from two PomA subunits. To clarify this and to better mimic the native environment of the stator unit, we reconstituted *Va*PomAB into membrane scaffold protein 1D1 (MSP1D1) nanodiscs, as well as full length, non-cleaved *Va*PomAB into saposin nanodiscs, with *E. coli* polar lipids, and determined the map resolution, at 3.9 Å and 6.3 Å, respectively (Extended Data Fig. S3-S4). In both cases, we were able to trace all the secondary structure elements of the PomAB complex, except those two PomA TM1 helices (Extended Data Fig. S5f, S5i). Comparison of the PomAB LMNG structure to the MSP1D1 lipid-reconstituted structure did not reveal major conformational differences (root-mean-square deviation of 0.36 Å) that could arise from detergent artifacts. This indicates that the flexibility of those two TM1 helices in the inactive stator unit is probably intrinsic, which might be functionally important during stator unit activation.

### Na^+^ ion binding sites and ion selectivity mechanism

Stator units use specific ions to power the flagellar motor rotation. Each MotB/PomB TM contains an aspartate (D24 in PomB) that is responsible for the binding and translocation of incoming ions from the periplasmic space to the cytoplasmic side (Fig. 2a). However, this aspartate is universally conserved among stator unit families (Extended Data Fig. S1b), obscuring the structural and mechanistic basis of the ion selectivity. The PomAB structure shows that D24 from two PomB chains sit in a different environment; D24 of Pom B chain 1 interacts with PomA, which we refer to as an engaged state; while D24 of PomB chain 2 points towards the cytoplasmic domain and breaks the interaction with PomA, which we refer to as a disengaged state (Fig. 3a). Examination of the high-resolution density map in the vicinities of these two aspartates reveals nonresidue densities. In site 1, close to the engaged PomB D24 (PomB chain1), the extra density is coordinated by oxygens from side chain hydroxyl groups of PomB T21 and D24, and backbone carbonyl groups of adjacent PomA P151 and PomB G20. A fifth coordinating interaction is made by a hydrogen bond from a water molecule near PomA A190, with the average distance between the center of the density and associated oxygens is 2.88 Å (Fig. 3b). In site 2, near the disengaged PomB D24, which is more flexible as indicated by the slightly blurred EM density of its acidic side chain, a globular density is well coordinated by oxygen atoms exclusively contributed by PomA TM3 and TM4: side chain hydroxyl groups of T158, T185 and T186, and exposed backbone carbonyl groups of G154 and A182, with an average distance between the density center and associated oxygen of 2.33 Å (Fig. 3c). Given the cation’s favorable local chemical environments in these two sites, and especially the typical geometry of Na^+^ coordination^48^ in site 2, we modeled these densities as Na^+^ ions, which were the most predominant cations in the protein purification buffer. To further validate the model, we performed two explicit solvent all-atom MD simulations (1 μs for each) and observed that the Na^+^ ion in site 1 was very stable, but the other Na^+^ ion in site 2 rapidly moved to an intermediate site formed by the side chain of D24, T158 and T186 and subsequentially to a location symmetric to site 1, and finally released to the cytoplasmic space (Extended Data Fig. S6c-d and Supplementary Movie 1). We also observed significant conformational dynamics of a few polar residues around site 2, especially T158, T186 in PomA chain 5 and D24 in PomB chain 2 (Extended Data Fig. S6a and S7). By contrast, T186 in PomA chain 2 and D24 in PomB chain 1 on the engaged site were however much more stable (Extended Data Fig. S6a-b and S7d).

**Fig. 3.**
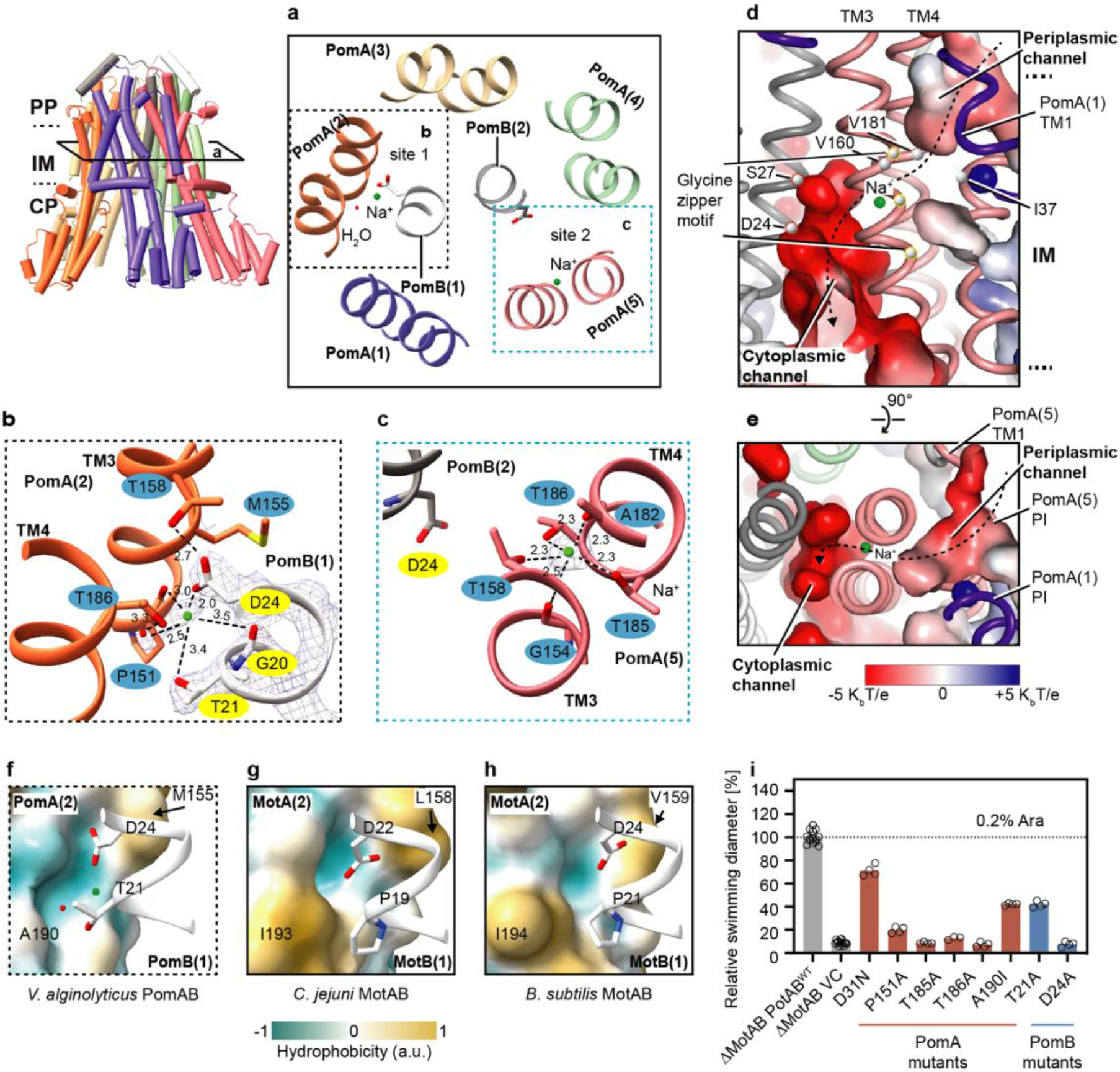
Ion binding sites, selectivity, and translocation pathway. **a**, Cross section view (corresponding to the view in left panel and rotated 90°) of Na^+^ ion binding sites (cyan spheres) in the vicinities of the two Asp24 from PomB. **b**, Details of the Na^+^ ion binding site near PomB(1) engaged Asp24. For clarity, corresponding EM densities are only overlapped in the region of PomB(1) Gly20-Asp24, Na^+^ ion, and water molecule. Hydrogen bonds are indicated as dashed lines with distances in angstroms. **c**, Details of the Na^+^ ion binding site near disengaged PomB(2) Asp24. EM density is overlaid on the Na^+^ ion. **d**, Na^+^ ion translocation pathway (dashed line with arrow). Periplasmic and cytoplasmic channels are indicated, with surface colored by electrostatic potential (positively charged, blue; negatively charged, red). Cα atoms of the residues forming the putative hydrophobic gate, of the glycines forming the glycine zipper motif, and of the PomB (2) S27 and D24 Cα are indicated and shown as spheres **e**, Top view of the Na^+^ ion translocation pathway. **f**, *Va*PomAB sodium ion binding environment near the engaged site. The surface of PomA is colored by hydrophobicity. **g**, Similar view as in **f**, but in the proton-driven stator unit *Cj*MotAB. **h**, Similar view as in **f**, but in the proton-driven stator unit *Bs*MotAB. **i**, Comparison of motility ability of the *Va*PotAB constructs and point mutants of the residues near the Na^+^ ion binding site or residues along Na^+^ translocation pathway.

The identification of the Na^+^ binding sites from EM density and the asymmetric conformational dynamics led us to speculate that at least part of the PomAB ion selectivity filter nests within the PomA subunit, and those three threonine residues (PomA T158, T185 and T186), which are conserved in all sodium-driven stator units (Extended Data Fig. S1a), account for the Na^+^ ion selectivity and transportation. Of note, the T158 from PomA chain 2, near the engaged PomB D24, does not directly contribute to the Na^+^ binding. Instead, it orients its side chain to establish a hydrogen bond with PomB D24 (Fig. 3b), indicating that a local conformational change occurs during Na^+^ ion transportation. Similarly, on the same site of the other three PomA subunits, we did not observe densities corresponding to a Na^+^ ion (Extended Data Fig. S8a-f), suggesting only one Na^+^ ion would be supplied during stator unit rotational steps.

To probe the critical role of key residues for the functional ion selectivity of Na^+^-driven stator unit, we first designed a chimeric PomAB (renamed as *Va*PotAB) by replacing PomB PGB with *S. enterica* MotB PGB, a strategy similar to that used in previous studies^49,50^. A plasmid encoding *Va*PotAB conferred a motile phenotype on soft agar plates when transformed into a mutant *Salmonella enterica* strain that lacks MotAB (Fig. 3i). We then made point mutations based on *Va*PotAB to evaluate the significance of the three key threonines on flagellar motor rotation by examining the motility phenotype. We found that substituting any of these three threonines to alanines abolishes bacterial motility, confirming the importance of these residues to stator unit function. The Na^+^ ion binding cavity therefore seems a strict requirement for ion selectivity. A K^+^ ion, which has a larger radius than Na^+^ ion (1.52 Å vs 1.16 Å) and has an average ligated bond distance of around 2.7-3.2 Å, cannot be accommodated in this cavity. On the other hand, H^+^ is too small to fill this cavity, and it is energy unfavorable for a H_3_O^+^ to be liganded with a coordination number of five. Therefore, K^+^ and H^+^ (or H_3_O^+^) cannot be used by PomAB as coupling ions. Divalent ions, such as Ca^2+^ and Mg^2+^, which would need further negatively charged residues to be neutralized and coordinated, are therefore not favored in this cavity either.

Additionally, we compared the PomAB structure with the available H^+^-driven stator unit structures, *C. jejuni* MotAB and *B. subtilis* MotAB, to explore the reason why H^+^-driven stator units cannot use sodium or other alkaline metals as coupling ions. In the part of the structure of the H^+^-driven stator unit that is equivalent to the corresponding Na^+^ binding site 2 in PomAB, two threonines (T158 and T185) are replaced by alanine, lacking oxygen in this cavity, likely precluding alkaline metal ion binding (Extended Data Fig. S1a). In the equivalent position of the PomB engaged D24, near the water molecule that coordinates the Na^+^ binding site 1, the H^+^-driven stator unit contains an isoleucine residue instead of an alanine or a polar residue, which makes this region hydrophobic and does not favor an alkaline metal ion coordination (Fig. 3f-h). Thus, both sites in the H^+^-driven stator unit lack the contact environment for alkaline metal ions, and these analyses further support the idea that the residue variability in PomA/MotA has a large influence on ion selectivity.

Analysis of the structure assembly interface between PomA and PomB subunits at the periplasmic level reveals that this inner contact interface is mainly lined by hydrophobic residues (Fig. 4a), with the thickness spanning around four helical turns (from PomB S27 to S38). It is therefore unlikely that an aqueous channel that mediates the Na^+^ ion flow through PomAB is formed in this region. Rather, a potential Na^+^ translocation pathway could be delineated based on the PomAB structure and our functional motility assay. It extends from the Na^+^ binding site 2 to the periplasmic space, delineated on one side by the PI helix and the beginning of TM2 helix from the same PomA subunit and on the other side by the end of TM1 from the adjacent PomA subunit (Fig. 3d-e). The ion translocation pathway in this part contains a hydrophobic gate (Fig. 3d), likely removing the hydration shell of the incoming Na^+^; and towards the periplasmic space, the translocation pathway is lined by several polar residues, such as D31, T33 and S34, and many of them are conserved (Extended Data Fig. S9b-c). The Na^+^ translocation pathway reaches to the PomB D24 and to PomA cytoplasmic domain inner lumen, where the surface electrostatic potential is very negative (Fig. 4d), and, together with the N-terminus of PomB that harbors several negatively charged residues (Extended Data Fig. S1b), might attract the incoming Na^+^. We also found that PomA TM3 contains a strictly conserved GXXGXXXG (residues G154-G161) motif, a typical ‘glycine zipper’ structure contributing to channel formation in many membrane proteins^51^. Glycines from the ‘glycine zipper’ motif face TM3 and TM4 assemble interface, holding the Na^+^ selectivity filter in a middle position, and together with the conserved P151, contributing to the main chain conformational elasticity of this region when a Na^+^ ion passes through TM3 and TM4 cleft (Fig. 3d, Extended Data Fig. S1a and S9c).

**Fig. 4.**
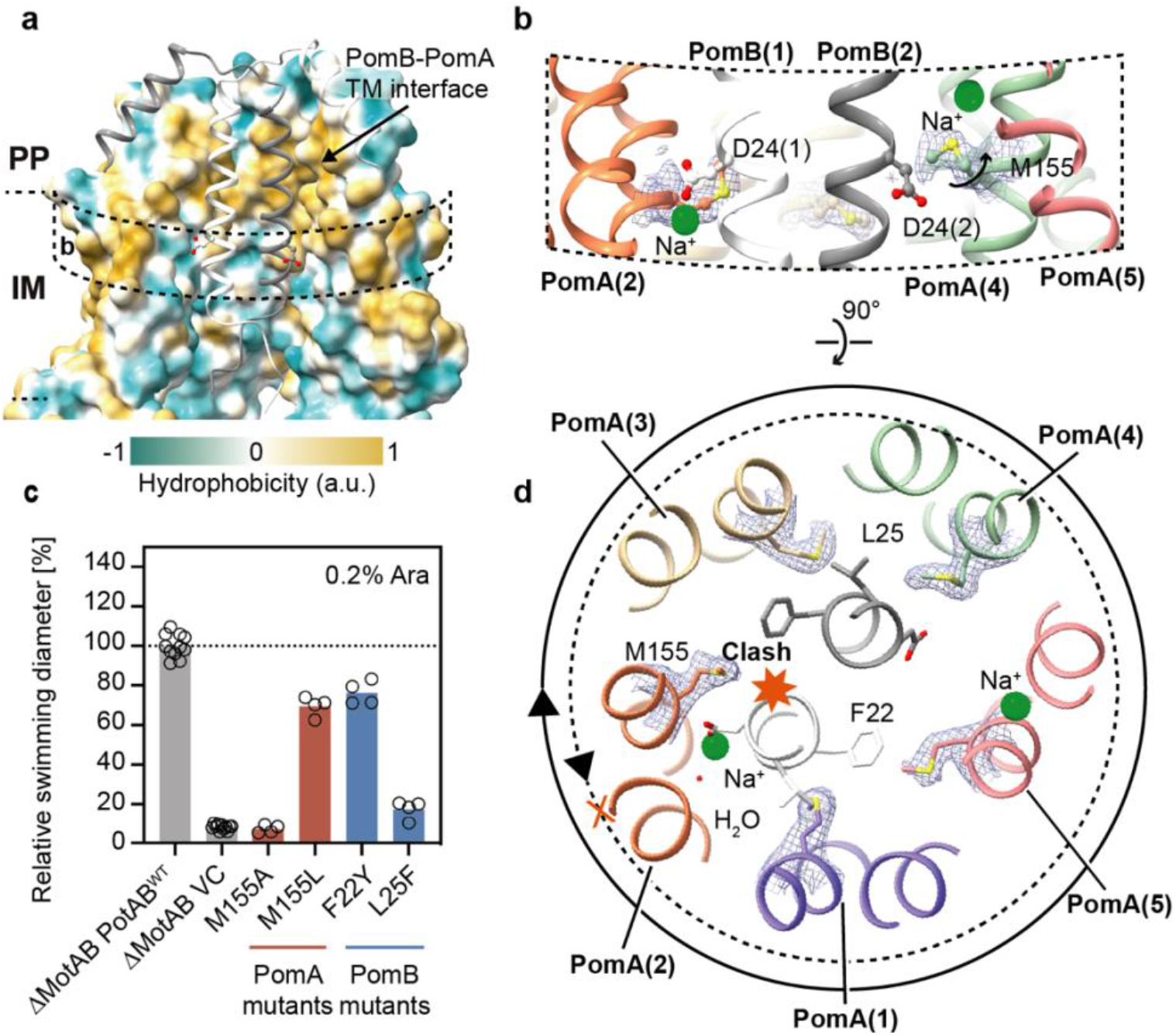
*Va*PomAB assembly interface and its directional rotation. **a**, *Va*PomAB assembly interface at the periplasmic space and transmembrane domain levels, with surface colored according to hydrophobicity. For clarity, the front two chains are deleted and PomB chains are shown as ribbon. **b**, Conformational isomers of M155 near PomB engaged D24 and disengaged D24. EM densities are overlaid on the side chain of M155. **c**, Comparison of motility ability of the *Va*PotAB constructs and point mutants of the residue M155, and residues from PomB near M155. **d**, Conformational isomers of M155 viewed from the top of the membrane. The solid circle indicates the rotational direction of PomA around PomB. A potential clash that would occur if PomA rotated CCW around PomB is indicated with a red heptagon.

From our explicit solvent MD simulations, we also observed that in the periplasmic side, the side chain of T33 was conformationally dynamic and surrounded by water molecules, which could occasionally diffuse to the space next to the side chain of T185 (Extended Data Fig. S7c and Supplementary Movie 2), therefore we propose that the hydration pocket form by T33 and a few other polar residues is the entry site of the proposed Na^+^ translocation pathway. Note that we did not observe a continuous hydration or Na^+^ translocation pathway to connect the periplasmic side and the Na^+^ site 2, probably because this structure was in the self-inhibited plugged state and the simulation time (1 μs) was also much shorter than the timescale of channel opening.

### The stator unit is primed for directional rotation

Having analyzed the ion selectivity mechanism of the stator unit family in the context of the high-resolution map of PomAB, we next sought to understand the structural basis of the rotational direction of the stator unit. Cryo-ET studies in *V. alginolyticus* and *Borrelia burgdorferi* basal bodies reveal that when the motor rotates in the CCW direction, its C-ring component FliG interacts with the stator unit cytoplasmic domain proximal side (the side facing the motor axis); while, when the motor is locked and rotates in the CW direction, FliG interacts with stator unit cytoplasmic domain distal side. The motor directional switching from CCW to CW rotation requires remodeling and expansion of the C-ring by changing its conformation upon receiving an intracellular chemotaxis signal^40,41^. Thus, the stator unit can drive both CW and CCW rotation of the flagellar motor with a relatively fixed position by anchoring itself to the peptidoglycan layer through the PGB motif.

Viewed from the plane of the inner membrane, we observe that the bulky hydrophobic side chain of M155 from PomA chain 2 is orientated horizontally to the engaged PomB D24 (Fig. 4b), revealing that M155 will sterically hinder PomA to CCW rotation around PomB at the engaged D24 site (Fig. 4d). Meanwhile, M155 from PomA chain 4 elevates its side chain to stride over the disengaged D24, for which the interaction with PomA is nearly absent, providing the required space for D24 to gather the Na^+^ ion from the selectivity cavity (Fig. 4b, 4d). We hypothesized that the bulky side chain of PomA position 155 is the stator unit directional rotation ‘reinforcement’ point. To test this hypothesis and verify the importance of the bulky side chain at this position, we first substituted this methionine residue with alanine. The M155A mutation abolished bacterial motility. In contrast, the replacement of methionine with leucine, a residue in the equivalent position often seen in H^+^-driven stator units, retained 80% motility. Increasing the size of the residues near PomA M155 from PomB (PomB F22Y and L25F) impaired motility (Fig. 4c). Therefore, our structural analysis and functional data confirm that a residue with a bulky side chain near the ion coupling site (D24 in PomB) is required to permit the correct rotation direction of the stator unit. Its conformational isomer (Extended Data Fig. S10), likely induced by the local structural rearrangement during the stator unit activation, is necessary to achieve flexibility in this region for ion transportation. This bulky hydrophobic residue is conserved not only in flagellar stator units, but also in other 5:2 rotary motors ^52^, suggesting a similar directional rotation ‘reinforcement’ mechanism (Extended Data Fig. S11). The stator unit is thus a preset CW rotary motor, which is tightly blocked by the trans mode conformation of the PomB plug motif at the periplasmic level before it incorporates onto the rotor. The geometry of the stator unit will not favor a model where PomA rotates CCW around PomB, when the ion motive force is reversed, due to the structural clashes (Fig. 4d) and negative electrostatic potential of PomA cytoplasmic inner lumen. This is consistent with early experiments showing that the stator unit is inactivated when the IMF is dissipated or reversed^53^, and that increased sodium concentration in the cytoplasm inhibits the rotation of PomAB^46^.

### PomA cytoplasmic domain and C-terminal helical motif

The stator unit cytoplasmic domain plays a crucial role during rotor incorporation, torque generation, and disassembly from the rotor^1,42^. The cytoplasmic domain of each PomA subunit contains four short helices that are almost vertical to the inner membrane. They peripherally surround the intracellular part of the PomA TM3 and TM4 helices and together form a compact helical bundle that protrudes approximately 35 Å into the cytosol (Fig. 5a-c). The cytoplasmic domains from five PomA subunits diverge towards their intracellular end, with the local resolution of this region decreasing considerably compared to the TMD. This is in line with the model B-factor distribution, where the PomAB cytoplasmic domain has a higher B-factor value (Extended Data Fig. S12), reflecting the flexibility of this region. The rotary stator unit generates torque by matching the complementary charged residues with the rotor FliG torque helix. This torque-generating mode is predicted to be conserved across bacterial species^24,26,54^. We divided the FliG torque helix-binding interface from the stator unit as follows: positively charged residues from one PomA subunit contribute to the principal face or (+) face, and negatively charged residues from the neighboring PomA subunit mainly contribute to the complementary face or (-) face (Fig. 5b-c). The PomAB structure allows us to map the locations of those key residues involved in stator rotor interaction. We found three positively charged residues from H1 and H4 at the (+) face, R88, K89, and R232, and two negatively charged resides from H2 and H3 at the (-) face, D114 and E96, that when the charge is suppressed or reversed, greatly impair motility (Fig. 5h). Importantly, the charge of R88 at the (+) face and D114 and E96 at the (-) face, whose side chains project toward the PomA intersubunit junction, are indispensable for motility, confirming that both (+) and (-) sides of PomA are necessary and directly involved in the interactions with FliG torque helix. Besides, R232 establishes an interdomain salt bridge with residue D85, and it is unlikely involved in the binding with FliG torque helix, rather, stabilizing helix bundle organization (Fig. 5b).

**Fig. 5.**
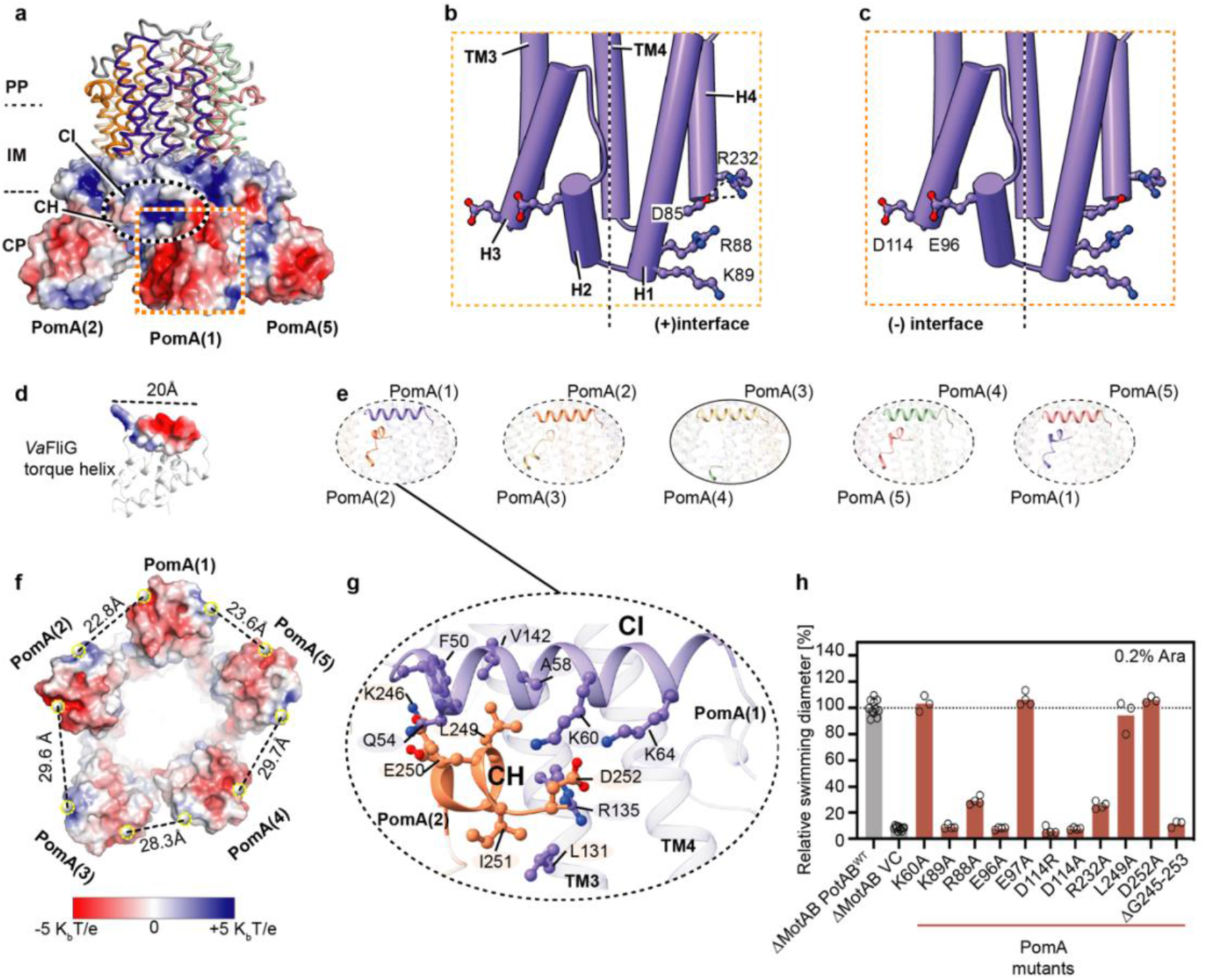
PomA cytoplasmic domain and C-terminal helical motif. **a**, PomAB cytoplasmic domain electrostatic potential. **b**, Locations of key residues responsible for FliG torque helix binding, highlighting the positively charged residues from the principal interface. **c**, Similar to **b**, but highlighting the negatively charged residues from the complementary interface. **d**, *Va*FliG C-terminal domain (based on homology modeling) containing the torque-generating helix is shown, and its length is indicated. **e**, Interactions between PomA CH helix and CI helix. One site without interaction is highlighted and circled with a solid line. **f**, Image from **a** viewed from the cytoplasmic domain. Distances between the center of mass of the residues K89, R88 and the center of mass of the residues D114, E96 from adjacent PomA subunits are given. **g**, Detailed interactions between CH motif and CI helix. Residues involved in interactions are shown as sticks. **h**, Comparison of motility ability of the *Va*PotAB constructs and point mutants of the residues involved in FliG torque helix interaction or PomA C-terminal truncation.

Unexpectedly, we found a helical (CH) motif right after the H4 helix in the PomA C-terminal part. The CH motif runs parallel to the membrane plane and attaches to the CI helix of a neighboring PomA subunit. In four PomA subunits, we could trace the entire CH motifs from residue K246 to its C-terminal end D253, with the contact between the CH and CI mainly mediated by electrostatic and hydrophobic interactions (Fig. 5g). The remaining CH motif is disordered, without any featured density observed (Fig. 5e). This disordered CH motif likely stems from the asymmetry of the PomAB assembly, where there are two PomA subunits on one side of PomB plug motifs and three on the other side and there is less space for this CH motif to interact with the neighboring PomA CI helix. The detachment of CH from CI at one intersubunit site results in the cytoplasmic domains of PomA forming an irregular pentagon, as shown by measuring distances of those charge residues responsible for FliG torque helix binding (the center of mass of K89 and R88 to the center of mass of D114 and E96) (Fig. 5f). The PomA C-terminal region is less conserved in length and sequence among stator unit subtypes (Extended Data Fig. S1b). We made a PomA C-terminal end truncation and found that PomA CH motif truncation completely abolished motility (Fig. 5h). Based on these findings and our structural analysis, we confirm that the PomA CH motif and the CH-CI interaction are critical to sustain stator unit function.

## Discussion

Since it was first revealed in a marine *Vibrio* species that its flagellar motor is driven by sodium-motive force^19,55^, the Na^+^-driven stator unit has been under intense functional and structural investigations for decades.

The trans conformation of the plug motifs seems to be a universal feature among the stator unit family and their structural configuration explains how this organization tightly restrain the rotation of the stator unit (Extended Data Fig. S13). The plug motifs also prevent ion influx into the cytoplasmic domain before the stator unit incorporates onto the rotor. Their distinct interaction environments caused by the imbalanced PomA_5_:PomB_2_ subunit stoichiometry also suggest their asymmetric release during the stator-rotor incorporation. The signal that promotes the periplasmic plug motif release is probably triggered by the cytoplasmic stator unit-rotor interaction upon the incorporation of the stator units into the motor, with the signal transmission route likely being through PomA transmembrane peripheral helices, particularly those two dynamic PomA TM1 helices. Plug motif release could then facilitate PomB PGB motifs dimerization, which can reach and anchor to the cell wall through recognition of the peptidoglycan components by the dimerized PGB interfacial grove, and this will produce a spatial tension preventing rebinding of the released plug motif to the activated stator unit. Therefore, only the rotor-incorporated unplugged stator units represent their fully activated states. Indeed, we were unable to purify the unplugged PomAB after deleting the PomB plug motif. Likely, the plug deletion PomAB complex did not assemble well and was toxic to the cells due to ion leakage, and the unplugged PomAB is more stable upon rotor incorporation.

The ion permeation pathway identified in the PomAB structure provides an energy advantage by shortening the sodium ion translocation path from the periplasmic side to the key ion-accepting residue PomB D24. PomB S27, a polar residue right above D24, may increase solvent accessibility (Fig. 3d). Additionally, the hydrophobic residues found at the periplasmic assembly interface of PomA and PomB may block the ion from flowing back to the periplasmic space, and they may also stabilize the stator unit by preventing it from falling apart during the stator unit’s dwell on the rotor (Fig. 4a). A recent study showed that when *E. coli* MotAB is replaced with an engineered PomAB (PomB PGB replaced with *E. coli* MotB PGB), at a low Na^+^ environment, the engineered PomAB can rapidly incorporate mutations, restoring the bacterial motility^56^ and reflect the adaptability of the stator unit. This is consistent with our results, where those mutations in the *Va*PotAB (PomB PGB replaced with *S. enterica* MotB PGB) granted the stator unit a gain-of-function phenotype in *S. enterica* (Extended Data Fig. S14a-b). Most of those mutation sites reside near the ion selectivity cavity (Extended Data Fig. S14c-d), including PomB G20, L28 and PomA L183, and upon mutation may modulate the ion specificity, probably enabling the stator unit to use both Na^+^ and H^+^ as coupling ions. Of note, in the H^+^-driven stator unit *C. jejuni* MotAB, the equivalent site of PomA L183 is phenylalanine (*Cj*MotA186), whose side chain adopts two conformations in the activated stator unit, affecting H^+^ translocation efficiency^35^. We also noticed that PomB L36Q has a gain-of-function phenotype. In the plugged PomAB structure, PomB chain1 L36 hydrophobically interacts with PomB chain 2 plug motif F47 (not PomB chain2 L36 with PomB chain 1 F47, due to asymmetric assembly) (Extended Data Fig. S8g-i). The L36Q mutation possibly decreases the plug motif binding energy and makes the stator unit more activable. Additionally, it is unlikely that PomB L36 lines the previously proposed ion translocation pathway in which it forms the dehydration gate with nearby hydrophobic residues^57,58^, as the L36A mutant has the same motility as the wild type phenotype (Extended Data Fig. S14a).

The observed CH-CI interactions and the detachment in one site as well as the irregular pentagonal shape of PomA cytoplasmic domain likely contribute to the process of stator unit assembly onto the rotor. We propose the following model for the dynamic stator unit binding to the rotor, in which the stator unit randomly orients towards the rotor and ‘measures’ the length of the FliG torque helix. Once both principal and complementary faces of the PomA cytoplasmic domain catch the FliG torque helix, possibly through (one of) the two shortest sites among five, which fit the length of FliG torque helix best (Fig. 5d, 5f), the stator unit is incorporated and is activated (Fig. 6a-c). This process could be assisted by FliL, a membrane protein recently shown to enhance the stator-rotor incorporation and stabilize the stator unit in its activated form^59,60^. During the activation, each PomA subunit near the disengaged PomB D24 supplies a Na^+^ from the ion selectivity cavity to couple the disengaged D24 with a Na^+^. Meanwhile, the engaged PomB D24 releases the coupled Na^+^ and together with M155 ensures CW rotation of PomA around PomB as viewed from outside of the membrane. At the same time, PomA cytoplasmic domain progressively interact with the FliG torque helix. The rotor will either be in CCW or CW rotation mode, depending on the conformation of the C-ring (Fig. 6d).

**Fig. 6.**
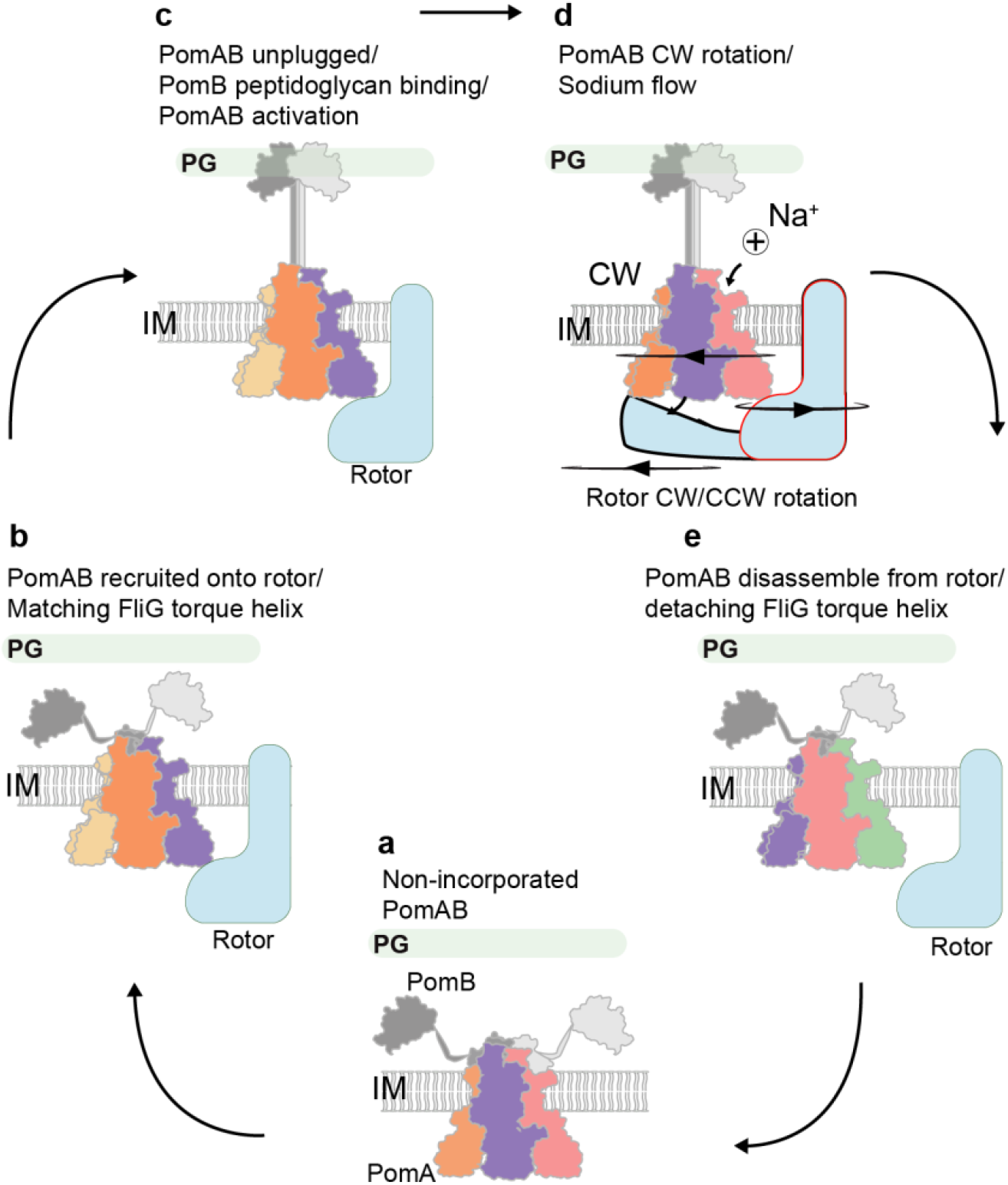
Models of PomAB activation and disassembly from rotor. **a**, An inactive stator unit is plugged autoinhibited. **b**, Inactive stator unit orients its cytoplasmic domain towards the rotor to contact FliG torque helix. **c**, The signal from the interaction between stator unit and rotor is transferred to the PomAB periplasmic domain, where it promotes the plug motifs release, followed by PomB PGB motifs dimerization and binding to the peptidoglycan layer. PomAB gets activated. **d**, In the activated PomAB, a sodium ion (represented by a sphere with a + symbol) passes through the PomA selectivity bind filter, and binds to PomB Asp24, triggering CW rotation of PomA around PomB. The rotor could rotate either CW or CCW direction, depending on how it interacts with the stator unit. **e**, Stator unit disassembly from the rotor when external torque is decreased.

Given the fact that stator units constantly assemble and disassemble around the rotor, depending on the requirement of external load, the asymmetric PomA cytoplasmic domain could also be advantageous for the deactivated stator unit to detach from the rotor. When the external load is decreased, which likely promotes the PGB motif of the PomB to disconnect from the cell wall, the plug motifs of the stator unit rebind to their inhibitory sites. This signal will transfer to the stator unit cytoplasmic domain, leading to its asymmetry and weakening the interactions between the stator unit and rotor, promoting the stator unit to separate from the rotor (Fig. 6e). The proposed model is reminiscent of the recently proposed ‘catch-bond’ mechanism, in which the interaction/bond becomes weaker under reduced force and is enhanced by rotation of the rotor^61,62^. However, the atomic structure of the whole flagellar motor with the assembled stator units is needed to fully understand the stator unit rotor incorporation mechanism and whether the asymmetric PomA cytoplasmic domain becomes symmetric during activation remains to be further investigated (Extended Data Fig. S15a-b).

In summary, we present the structures of *Va*PomAB in both detergent and lipidic environments. The cryo-EM maps not only provide a detailed structure assembly of the Na^+^-driven stator unit, but also enable us to assign the ion binding sites, which in turn allows us to address the enigmatic mechanism of stator unit ion selectivity. Our structural analysis and functional experiments support that the stator unit is a CW unidirectional rotary motor and this is achieved by a hydrophobic directional rotation ‘reinforcement’ point. The PomB plug motifs organization and discovery of PomA C-terminal helical motif further expand our view about the stator unit activation and rotor incorporation.

## Materials and methods

### *Va*PomAB purification with LMNG detergent

The DNA sequence coding for *Va*PomAB was amplified from *Vibrio alginolyticus* (ATCC 17749) and subcloned into a modified pET vector containing a C-terminal twin-Strep-tag. A human rhinovirus (HRV) 3C protease cleavage site (GTLEVLFQGPGGS) was inserted between the PomB plug motif and the peptidoglycan binding domain (between residues Gln95 and Gln96). PomAB complex was expressed in *E. coli* Overexpress™ C43(DE3) cells (LuBioScience GmbH). Cells were cultured in 8 l LB medium supplemented with 50 μg/ml ampicillin at 37°C, and protein expression was induced with 0.5 mM IPTG at OD_600_ 0.6. Cells were incubated for another 16 hours at 20°C before harvesting. The cell pellet was resuspended in buffer A (20 mM HEPES pH 7.5, 300 mM NaCl) with 30 μg/ml of DNase I and 50 μg/ml of lysozyme and incubated at 4°C for 30 min before passaging it through an EmulsiFlex-C5 homogenizer at 15,000-20,000 pound-force per square inch. Unbroken cells were removed by centrifugation at 8000 rpm for 15 min. Membranes were then sedimented at 41,000 rpm for 1 hour and stored at −20°C after flash freezing with liquid nitrogen.

For protein purification, membranes were solubilized in buffer A supplemented with 2% (w/v) Lauryl Maltose Neopentyl Glycol (LMNG), 10% glycerol, and protease inhibitors (protease inhibitor cocktail tablets, EDTA-free, Roche Diagnostics GmbH) for 2 hours at 4°C while shaking on a rocking platform, and then ultracentrifuged for 30 min at 28,000 rpm. The supernatant was added to a gravity flow column containing 2 ml Strep-Tactin® Superflow® resin (IBA) pre-equilibrated with washing buffer (buffer A with 10% glycerol and 0.005% LMNG). Resins were washed five times with 4 column volumes of washing buffer and Strep tagged protein was eluted with elution buffer (Buffer A, 10% glycerol, 0.005% LMNG and 10 mM desthiobiotin). The protein complex was then concentrated until reaching a volume of 0.5 ml. HRV-3C protease was added to the *Va*PomAB sample, with a protein:protease ratio of 5:1 (w/w) and incubated at 4°C overnight. The sample was loaded onto a Superose® 6 Increase 10/300 GL (Merck) column, pre-equilibrated with buffer A with 0.002% LMNG. The peak fractions corresponding to the protein complex were concentrated to about 16-20 mg/ml using a centrifugal filter with a PES membrane (Sartorius) and used for preparation of cryo-EM sample grids immediately.

### *Va*PomAB MSP1D1 and Saposin lipid nanodisc reconstitution

To reconstitute *Va*PomAB into lipid nanodiscs with MSP1D1, 500 μl of 2 mg/ml purified *Va*PomAB without PomB PGB was mixed with *E. coli* polar lipids and MSP1D1 in a molar ratio of 1:156:6.25 (VaPomAB:lipids:MSP1D1). The reaction was incubated at 4°C with mild agitation for 5 min. Bio-beads (300 mg per ml reaction) were added and incubated overnight to remove the detergent. Bio-beads were filtered out the next day using a PVDF 0.22 μm Centrifugal Filter (Durapore) tube. The sample was then injected into a Superose® 6 Increase 10/300 GL (Merck) column, which was pre-equilibrated with buffer A. The peak fractions corresponding to the protein complex in lipid nanodiscs of MSP1D1 were pooled, concentrated and used for cryo-EM grids preparation.

To reconstitute *Va*PomAB into lipid nanodiscs with saposin, 300 μl of 6 mg/ml full length purified *Va*PomAB (without protease insertion) was mixed with *E. coli* polar lipids (10 mM; 200 μl) and incubated at room temperature for 10 min. Saposin (6.7 mg/ml; 350 μl) was added into the reaction and incubated for 2 min. The molar ratio of PomAB, lipids and saposin was 1:300:35, respectively. The reaction was diluted with 2 ml buffer A to initiate the reconstitution and incubated on ice for an additional 30 min. 700 mg of bio-beads were added and incubated overnight to remove the detergent. The rest of the steps were the same as when *Va*PomAB was reconstituted into MSP1D1 nanodiscs.

### Cryo-EM grids preparation and cryo-EM data collection

To break the preferential particle orientation, 0.0125% CHAPSO (final concentration) was added into the sample before grid preparation. 2.7 μl of freshly purified sample was applied onto glow-discharged (30 s, 5 mA) grids (Quantifoil R 0.6/1 300 mesh Cu or Ultrafoil 0.6/1 300 mesh Au) and plunge-frozen into liquid ethane using a Vitrobot Mark IV (FEI, Thermo Fisher Scientific) with the following parameters: 4°C, 100% humidity, 7 s wait time, 4-4.5 s blot time, and a blot force of 25. Movies were collected using the semi-automated acquisition program EPU (FEI, Thermo Fisher Scientific) on a Titan Krios G2 microscope operated at 300 keV paired with a Falcon 3EC direct electron detector (FEI, Thermo Fisher Scientific). Images were recorded in an electron counting mode, at 96,000x magnification with a calibrated pixel size of 0.832 Å and defocus range of 0.8 to 3 μM. For the *Va*PomAB sample purified in LMNG, 6,467 micrographs were collected, with each micrograph containing 40 frames and a total exposure dose of 37.98 (e/Å^2^). For the *Va*PomAB sample reconstituted into saposin nanodiscs, 3,927 micrographs were collected, with each micrograph containing 40 frames and a total exposure dose of 37 (e/Å^2^). For the *Va*PomAB MSP1D1 sample, 5,450 micrographs were collected, with each micrograph containing 40 frames and a total exposure dose of 40 (e/Å^2^).

### Image processing

To keep the image data processing consistent, all the datasets were processed using cryoSPARC version 3.3.2, unless otherwhere stated. Patch motion correction was used to estimate and correct frame motion and sample deformation (local motion). Patch Contrast function (CTF) estimation was used to fit local CTF to micrographs. Micrographs were manually curated to remove the bad ones (relatively ice thickness thicker than 1.05 and CTF value worse than 3.2 Å for LMNG dataset; relatively ice thickness thicker than 1.1 and CTF value worse than 5 Å for MSP1D1 nanodisc dataset; relatively ice thickness thicker than 1.2 and CTF value worse than 5 Å for Saposin nanodisc dataset). Particles were picked using the Topaz software implemented in cryoSPARC ^63^. Basically, Topaz extract was used with a pre-trained model with a pre-tested particle threshold value. Particles were extracted with a box size of 400 pixels and Fourier crop to box size of 100 pixels. Duplicated particles were removed using a minimum separation distance criteria of 60 Å, which means that the distance between the centers of two neighboring particles should be larger than 60 Å. One round of 2D classification was then performed, followed by ab-initio reconstruction. Heterogeneous refinement was used to get rid of the junk particles. Particles were re-extracted with full box size (400 pixels). Non-uniform refinement was applied with a dynamic mask to obtain a high-resolution map. Local refinement was additionally performed with a soft mask surrounding *Va*PomAB complex in order to achieve a higher resolution map. The number of micrographs, total exposure values, number of particles used for final refinement, and map resolution values for all datasets are summarized in Table S1.

### Atomic model building, refinement, and validation

ColabFold ^64^ was used to predict the structure of PomA pentamer ^65^ and manually fit the model into the density by using UCSF ChimeraX ^66^. The model was refined in Coot ^67^, and PomB TM and plug motif was manually modelled. The model was then refined against the map using PHENIX real space refinement ^68^.

### Molecular dynamics simulation of PomAB

The system was constructed by embedding the cryo-EM structure of PomAB into a flat, mixed lipid bilayer consisting of 16:0-18:1 phosphatidylethanolamine (POPE) and 1-palmitoyl-2-oleoyl phosphatidylglycerol (POPG) at a 4:1 ratio using the Membrane Builder tool of CHARMM-GUI webserver ^69^. Explicit water was added using the TIP3P water model, and the system charge was neutralized with sodium ions and solvated in a cubic water box containing 0.15 M NaCl. The size of the box was 11.0 nm, 11.0 nm, and 11.5 nm in the x, y and z dimension, respectively, resulting in ∼144,000 atoms in total. The CHARMM36m force field ^70^ was used for the protein, and the CHARMM36 lipid force field ^71^ was used for all lipid molecules. Note that the WYF correction was included in the force field to improve the description of the cation-π interactions ^72^. The temperature was kept constant at 310 K using the V-rescale algorithm with a 2 ps coupling constant, and the pressure at 1.0 bar using the Parrinello-Rahman barostat ^73^ with a 5 ps time coupling constant. A cutoff of 1.2 nm was applied for the van der Waals interactions using a switch function starting at 1.0 nm. The cutoff for the short-range electrostatic interactions was also at 1.2 nm and the long-range electrostatic interactions were calculated by means of the particle mesh Ewald decomposition algorithm with a 0.12 nm mesh spacing. A reciprocal grid of 96 × 96 × 96 cells was used with 4th order B-spline interpolation. MD simulations were performed using Gromacs2021.5 ^74^. Two independent simulations were performed, each for one μs. Analysis of the MD trajectories was performed using the Gromacs gmx and GROmaρs tools ^75^.

### Bacterial strains and growth

*Escherichia coli* and *Salmonella enterica* serovar *Typhimurium* LT2 (J. Roth) (ATCC 700720) were grown at 37°C with aeration at 180 rpm in lysogeny broth (LB medium) [10 g/l tryptone, 5 g/l yeast extract and 5 g/l NaCl]. For solid agar plates, 1.5% (w/v) of agar-agar was added, alternatively to test swimming motility 0.3% (w/v) of agar-agar was supplemented. All strains used in this study are listed in the supplement information Table S2. For strains harboring a plasmid carrying a resistance marker selected media were supplemented with chloramphenicol (12.5 μg/ml). Induction experiments were performed in the presence of arabinose (0.2%).

### DNA manipulation

Plasmids were constructed according to standard cloning techniques as described elsewhere (ISBN 0879695773). In brief, rolling circle, around the horn PCR and overlap PCR were applied to generate point mutations in *pomA* or *pomB*, respectively. The primers used in this study are listed in the supplement information Table S3. For DNA amplification Q5 polymerase was used and for verification OneTaq polymerase (both purchased from NEB, Ipswich, MA, USA). All plasmids were verified by sequencing.

### Motility assay

To assess the swimming motility of *Va*PotAB mutants respective strains were inoculated in LB medium supplemented with chloramphenicol. From overnight cultures, soft agar plates containing the selective marker and supplemented with or without arabinose were inoculated with 2 μl and incubated at 37°C. Once a decent halo was visible, plates were scanned. From these pictures swimming diameters were evaluated using Fiji (10.1038/nmeth.2019).

### Figure preparation

Figures were prepared using ChimeraX ^66^, PyMOL, GraphPad Prisim 9 and Adobe Illustrator. Surface buried area and solvation free energy was calculated using the online webserver PDBePISA ^76^.

## Data and Code Availability

Atomic coordinates for VaPomAB in LMNG detergent and VaPomAB in MSP1D1 nanodisc were deposited in the Protein Data Bank under accession codes PDB: 8BRD and 8BRI, respectively. The corresponding electrostatic potential maps were deposited in the Electron Microscopy Data Bank (EMDB) under accession codes EMDB: EMD-16212and EMD-16215, respectively. The electrostatic potential map for full length VaPomAB in Saposin nanodisc was deposited in the EMDB under accession code EMDB: EMD-16214.

## Author contribution

N.M.I.T. supervised the project and acquired funding. H.H. expressed, purified, optimized, prepared cryo-EM grids, collected cryo-EM data, and determined the structure of *Va*PomAB and the structures of *Va*PomAB in nanodiscs. M.S. helped with protein expression, purification, and cryo-EM grid preparation at the beginning of this project. P.F.P. did the motility assay and interpreted data together with M.E.. W.Y. and Z.L. performed the molecular dynamics simulations. M.S., A.R.-E., ad Y.M.Y. helped with data analysis and figure preparation. N.W helped with data interpretation. H.H. built and refined the structure models, prepared figures and wrote the first draft of the manuscript with input from all the authors, which was then edited by N.M.I.T. and M.E.. All authors contributed to the revision of the manuscript.

## Acknowledgements

The Novo Nordisk Foundation Center for Protein Research is supported financially by the Novo Nordisk Foundation (NNF14CC0001). N.M.I.T. acknowledges support from DFF grant (8123-00002B) and NNF Hallas-Møller Emerging Investigator grant (NNF17OC0031006). M.E. acknowledges support by the European Research Council (ERC) under the European Union’s Horizon 2020 research and innovation program (agreement no. 864971). Y.W. acknowledges the financial support from National Key Research and Development Program of China (No. 2021YFF1200404) and the Fundamental Research Funds for the Central Universities of China (No. K20220228), as well as the access to computational resources from the Information Technology Center and State Key Lab of CAD&CG, ZheJiang University. H.H. acknowledges support from Lundbeck Foundation postdoc R347-2020-2429. We thank the Danish Cryo-EM Facility at the Core Facility for Integrated Microscopy (CFIM) at the University of Copenhagen and Tillmann Pape and Nicholas Heelund Sofos for support during data collection.

## Figure legends

**Fig. S1 Protein sequence alignment of *Va*PomA and *Va*PomB homologs from different bacterial species**.

**a-b**, Multiple-sequence alignment of PomA (**a**) and PomB (**b**). The proteins are grouped into two families: sodium- and proton-driven stator units. In the case of *Cs*MotAB, whose cryo-EM structure is available, the ion type is ambiguous, and therefore it is labeled with a question mark. *Va*PomAB residue numbers (in red) are given above the sequences. Helices are indicated by solid boxes. Residues that are identical or partially conserved are highlighted in red and orange, respectively. Residues that are critical for sodium ion selectivity in PomAB (T158, T185 and T186) are marked with a star. Dashed line above the PomB sequence indicates that the structure was not resolved in the PomAB complex cryo-EM map. PomB PGB domain is also indicated above the sequence alignment. PomA C-terminal helical motif is highlighted by a semi-transparent green box. Sequences aligned: *Vibrio alginolyticus Va*PomAB; *Vibrio mimicus Vm*PomAB; *Shewanella oneidensis So*PomA *and So*PomB; *Bacillus pseudofirmus Bp*MotPS; *Bacillus subtilis Bs*MotPS, *Bs*MotAB; *Bacillus alcalophilus Ba*MotPS; *Escherichia coli Ec*MotAB; *Salmonella enterica Se*MotAB; *Campylobacter jejuni Cj*MotAB; *Clostridium sporogenes Cs*MotAB.

**Fig. S2 Cryo-EM of *Va*PomAB in LMNG detergent**.

**a**, A representative SEC profile of LMNG detergent purified *Va*PomAB complex. The fraction used for preparing cryo-EM grids is indicated with a pink rectangular bar. **b**, SDS gel from **a** is shown. **c-d**, Flowchart of the data processing of *Va*PomAB in LMNG in cryoSPARC that results in the final cryo-EM structure of *Va*PomAB at around 2.5 Å resolution after non-uniform refinement. **e**, Gold standard (0.143) Fourier shell correlation (GSFSC) curves for *Va*PomAB in LMNG. **f**, Particle directional distribution of *Va*PomAB in LMNG. **g**, Cryo-EM density map of *Va*PomAB in LMNG detergent colored by local resolution (in Å) estimated in cryoSPARC. **h**, Representative model segments fitted into EM density.

**Fig. S3 Cryo-EM of *Va*PomAB in MSP1D1 lipid nanodisc**.

**a**, SDS gel analysis of purified *Va*PomAB in MSP1D1 lipid nanodisc. **b**, Flowchart of the data processing of *Va*PomAB in MSP1D1 lipid nanodisc in cryoSPARC that results in the final cryo-EM structure. **c**, The final cryo-EM map of *Va*PomAB in MSP1D1 lipid nanodisc at around 3.9 Å resolution. **d**, Cryo-EM density map of *Va*PomAB in MSP1D1 lipid nanodisc colored by local resolution (in Å) estimated in cryoSPARC. **e**, Gold standard (0.143) Fourier shell correlation (GSFSC) curves for *Va*PomAB in MSP1D1 lipid nanodisc. **f**, Particle directional distribution of *Va*PomAB in MSP1D1nanodisc. **g**, Representative model segments fitted into EM density.

**Fig. S4 Cryo-EM of full length *Va*PomAB in saposin lipid nanodisc**.

**a**, SDS gel analysis of the purified full length *Va*PomAB in saposin lipid nanodisc. **b**-**c**, Flowchart of the data processing of full length *Va*PomAB in saposin lipid nanodisc in cryoSPARC that results in the final cryo-EM structure. **d**, The final cryo-EM map of *Va*PomAB in saposin lipid nanodisc at around 6.3 Å resolution after local refinement. **e**, Gold standard (0.143) Fourier shell correlation (GSFSC) curves for *Va*PomAB in saposin lipid nanodisc. **f**, Particle directional distribution. **g**, Cryo-EM density map of *Va*PomAB in saposin lipid nanodisc colored by local resolution (in Å) estimated in cryoSPARC.

**Fig. S5 Dynamics of *Va*PomA PI and TM1 helices**.

**a-c**, Representation of the *Va*PomAB LMNG unsharpened electrostatic potential maps at low threshold showing the conformational dynamic of PI helices that interact with PomB plug motifs, and the flexibility of the corresponding TM1 helices. **d**-**f**, Representation of the *Va*PomAB MSP1D1 lipid nanodisc unsharpened electrostatic potential maps at low threshold. **g**-**i**, Representation of the full length *Va*PomAB saposin lipid nanodisc unsharpened electrostatic potential maps at low threshold.

**Fig. S6 Na**^**+**^ **translocation pathway and dynamics of PomB D24**.

**a**-**b**, The trajectories of the side chain dynamics of D24 in PomB chain 1 and 2 obtained from two independent MD simulations. **c**, The cryo-EM Na^+^ binding sites. The modelled Na^+^ ions are shown by blue spheres. **d**, The Na^+^ binding sites captured in MD simulations. The average density of Na^+^ ions is represented by red mesh in **c** and **d**.

**Fig. S7 Hydration of T33 and the Na**^**+**^ **translocation pathway and side chain dynamics of T158, T185 and T186 obtained from explicit solvent MD simulations**.

**a**-**b**, The hydration and Na^+^ binding in the engaged and disengaged state, respectively. The average density of water molecules is represented by mesh in green. **c**, A snapshot from the MD simulations to show the hydration of T33 in PomA chain 5. **d**-**f**, The MD trajectories of the side chain dynamics of T186, T185 and T158 in PomA chain 2 and 5.

**Fig. S8 densities of ion selectivity cavities**.

**a**, View from the plane of the membrane, showing the position of ion selectivity cavity within the complex. **b**-**f**, ion selectivity cavities from PomA chains 1 to 5. EM densities are overlaid on the corresponding local regions. **g**-**i**, L36 from PomB chain 1 and chain 2 interaction environments, showing that PomB chain 1 L36 interacts PomB chain 2 F47.

**Fig. S9 Conservation (calculated with ConSurf) analysis of *Va*PomA and *Va*PomB**.

**a**-**b**, Conservation (calculated with ConSurf) of the surface residues of *Va*PomA from external and internal sides; Cα atom representation (shown as spheres) of the model colored by conservation. **c**, Conservation of the residues of the Na^+^ ion selectivity filter and permeation pathway from the periplasmic side, both external and internal views are shown. **d**, Conservation of the residues of PomA cytoplasmic domain, highlighting the locations of the positively charged residues from the principal face involved in FliG torque helix binding. **e**, Same as in **d**, but highlighting negatively charged residues from the complementary face. **f**, Conservation of the surface residues of *Va*PomB, highlighting the strictly conserved residues. **g**, Same as in **f**, but rotated 180 degrees.

**Fig. S10 Conformational isomers of *Va*PomAB M155**.

**a**, View from the plane of the membrane, showing the position of PomA M155 within the complex. **b**-**f**, M155 isomers from PomA chains 1 to 5. EM densities are overlaid on the side chains of M155. **g**, Conformational isomers of M155 viewed from the top of the membrane.

**Fig. S11 5:2 rotary motor directional rotation ‘reinforcement’ point**.

**a**, Proton-driven flagellar stator unit *Cj*MotAB (PDB: 6YKM). **b**, Conformational isomers of L158 near MotB engaged D24 and disengaged D24. **c**, Conformational isomers of L158 viewed from the top of the membrane. Solid circle indicates the rotational direction of MotA around MotB. The potential clash that would occur if PomA rotated CCW around PomB is indicated with a red heptagon. **d**, Proton-driven Ton ExbB-ExbD complex (PDB: 6TKI). **e**, Conformational isomers of L145 near ExbD engaged D25 and disengaged D25. **f**, Conformational isomers of ExbB L145 viewed from the top of the membrane. Solid circle indicates the rotational direction of ExbB around ExbD. The potential clash that would occur if ExbB rotated CCW around ExbD is indicated with a red heptagon.

**Fig. S12 *Va*PomAB model B-factor distribution**.

Top (**a**) and side views (**b**) of the PomAB model (LMNG dataset) colored by B-factor distribution (atomic displacement factor).

**Fig. S13 H**^**+**^**- and Na**^**+**^**-driven stator units PomB/MotB plug motifs organization**.

**a**, Side view of the proton-driven stator unit *Cj*MotAB in its auto-inhibited state.**b**, *Cj*MotAB viewed from the top of the membrane. **c**, Side view of the sodium-driven stator *Va*PomAB in its auto-inhibited state. **d**, *Va*PomAB viewed from the top of the membrane. Rotational direction of the stator unit is indicated. **e**, The unique trans mode organization of the plug motifs tightly blocks the CW rotation of the stator unit.

**Fig. S14 Mutational analysis for *Va*PomA and *Va*PomB plotted onto the *Va*PomAB structure**.

**a**-**b**, The motility phenotypes of VaPotAB PomA (**a**) and PotB (**b**) point mutants were analyzed using soft-agar motility plates containing 0.2% agar. **c**-**d**, Swimming efficiency of the *Va*PotAB point mutants, showing the mutated residues as Cα spheres on the PomA (purple) and PomB (white) structure.

**Fig. S15 Conformational changes of PomA cytoplasmic domain during stator unit activation and disassembly from the rotor**.

**a**, PomA cytoplasmic domain is asymmetric, and one site of the CH-CI detachment is indicated in dashed line. Inactive stator unit orients its cytoplasmic domain towards the rotor to contact FliG torque helix through FliG torque helix ‘matching sites’ ((1)-(2)). During the activation, all five CH-CI interactions established, and PomA cytoplasmic domain becomes symmetric ((3)-(4)). The rotor could rotate either CW or CCW direction, depending on how it interacts with the stator unit. Stator unit disassembly from the rotor when external torque is decreased ((5)-(6)). **b**, In this model, during the stator unit activation, PomA cytoplasmic domain remains asymmetric ((3)-(4)); one site of the CI helix attaches to the PI helix and the adjacent CI helix detaches from the PI helix, sequentially creating a FliG torque helix ‘catching’ site that interacts with the FliG torque helix.

## Notes

### Competing Interest Statement

The authors have declared no competing interest.

