## Supplemental figures for "Mechanisms of ion selectivity and rotor coupling in the bacterial flagellar sodium-driven stator unit"

a

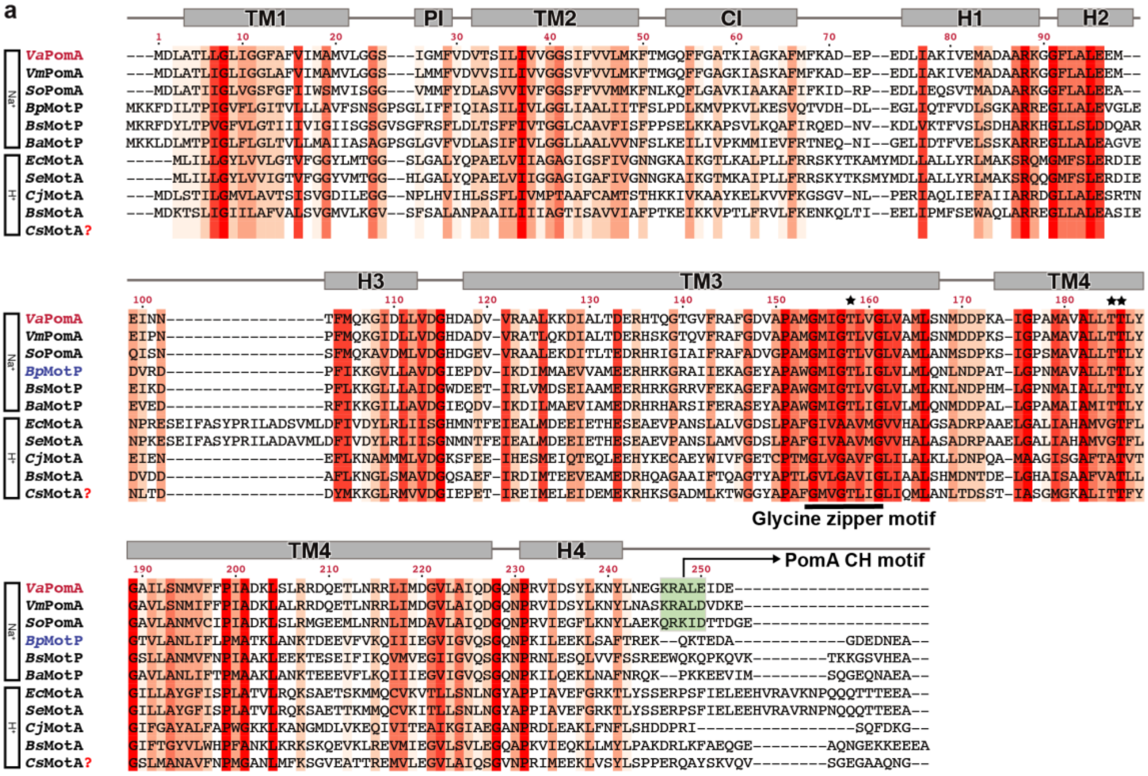

TM1
PI
TM2
CI
H1
H2
H3
TM3
TM4
H4
VaPomA
VmPomA
SoPomA
BpMotP
BsMotP
BaMotP
EcMotA
SeMotA
CjMotA
BsMotA
CsMotA?
Na+
H+
Glycine zipper motif
PomA CH motif


b

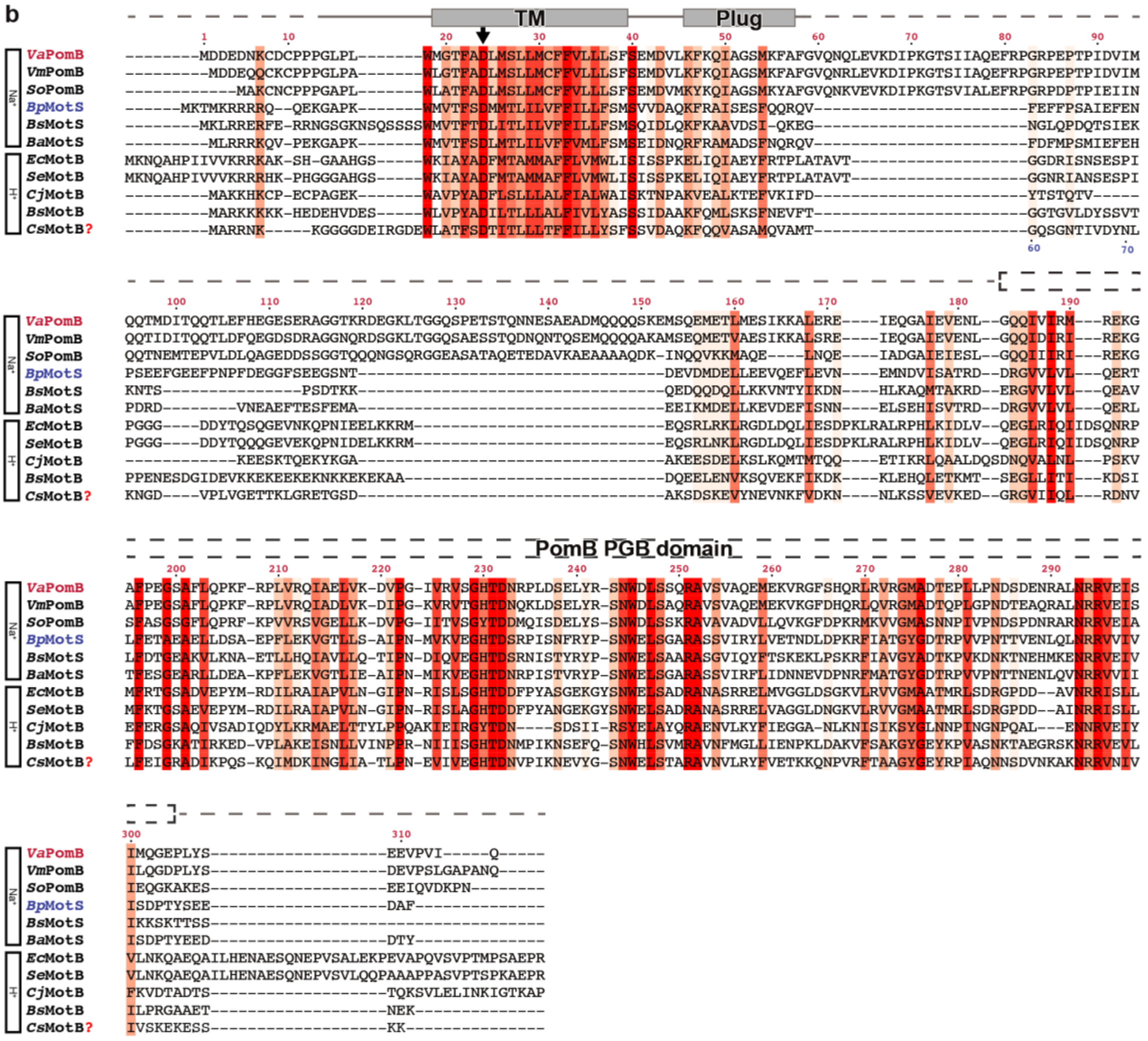

TM
Plug
PomB PGB domain
VaPomB
VmPomB
SoPomB
BpMotS
BsMotS
BaMotS
EcMotB
SeMotB
CjMotB
BsMotB
CsMotB?
Na+
H+

**Fig. S1 Protein sequence alignment of *Va*PomA and *Va*PomB homologs from different bacterial species.**

**a**-**b**, Multiple-sequence alignment of PomA (**a**) and PomB (**b**). The proteins are grouped into two families: sodium- and proton- driven stator units. In the case of *Cs*MotAB, whose cryo-EM structure is available, the ion type is ambiguous, and therefore it is labeled with a question mark. *Va*PomAB residue numbers (in red) are given above the sequences. Helices are indicated by solid boxes. Residues that are identical or partially conserved are highlighted in red and orange, respectively. Residues that are critical for sodium ion selectivity in PomAB (T158, T185 and T186) are marked with a star. Dashed line above the PomB sequence indicates that the structure was not resolved in the PomAB complex cryo-EM map. PomB PGB domain is also indicated above the sequence alignment. PomA C-terminal helical motif is highlighted by a semi-transparent green box. Sequences aligned: *Vibrio alginolyticus Va*PomAB; *Vibrio mimicus Vm*PomAB; *Shewanella oneidensis So*PomA *and So*PomB; *Bacillus pseudofirmus Bp*MotPS; *Bacillus subtilis Bs*MotPS, *Bs*MotAB; *Bacillus alcalophilus Ba*MotPS; *Escherichia coli Ec*MotAB; *Salmonella enterica Se*MotAB; *Campylobacter jejuni Cj*MotAB; *Clostridium sporogenes Cs*MotAB.

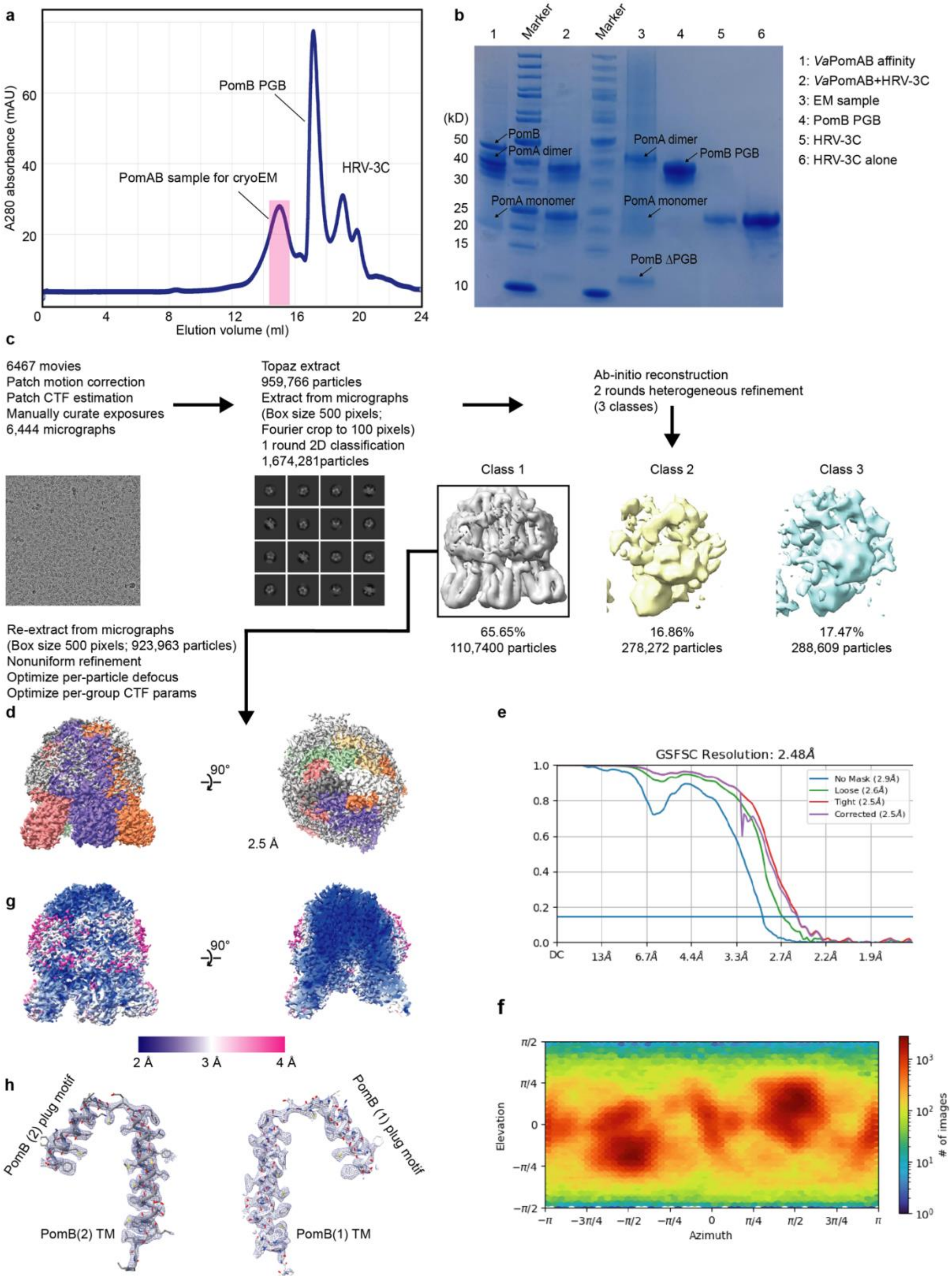
a
A280 absorbance (mAU)
Elution volume (ml)
PomB PGB
HRV-3C
PomAB sample for cryoEM
b
Marker
1: *Va*PomAB affinity
2: *Va*PomAB+HRV-3C
3: EM sample
4: PomB PGB
5: HRV-3C
6: HRV-3C alone
(kD)
PomB
PomA dimer
PomA monomer
PomB PGB
PomB ΔPGB
c
6467 movies
Patch motion correction
Patch CTF estimation
Manually curate exposures
6,444 micrographs
Topaz extract
959,766 particles
Extract from micrographs
(Box size 500 pixels;
Fourier crop to 100 pixels)
1 round 2D classification
1,674,281particles
Ab-initio reconstruction
2 rounds heterogeneous refinement
(3 classes)
Class 1
Class 2
Class 3
65.65%
110,7400 particles
16.86%
278,272 particles
17.47%
288,609 particles
Re-extract from micrographs
(Box size 500 pixels; 923,963 particles)
Nonuniform refinement
Optimize per-particle defocus
Optimize per-group CTF params
d
90°
2.5 Å
e
GSFSC Resolution: 2.48Å
No Mask (2.9Å)
Loose (2.6Å)
Tight (2.5Å)
Corrected (2.5Å)
DC
13Å
6.7Å
4.4Å
3.3Å
2.7Å
2.2Å
1.9Å
g
90°
2 Å
3 Å
4 Å
f
Elevation
Azimuth
### of images
h
PomB (2) plug motif
PomB (1) plug motif
PomB(2) TM
PomB(1) TM

**Fig. S2 Cryo-EM of *Va*PomAB in LMNG detergent.**

**a**, A representative SEC profile of LMNG detergent purified *Va*PomAB complex. The fraction used for preparing cryo-EM grids is indicated with a pink rectangular bar. **b**, SDS gel from **a** is shown. **c-d**, Flowchart of the data processing of *Va*PomAB in LMNG in cryoSPARC that results in the final cryo-EM structure of *Va*PomAB at around 2.5 Å resolution after non-uniform refinement. **e**, Gold standard (0.143) Fourier shell correlation (GSFSC) curves for *Va*PomAB in LMNG. **f**, Particle directional distribution of *Va*PomAB in LMNG. **g**, Cryo-EM density map of *Va*PomAB in LMNG detergent colored by local resolution (in Å) estimated in cryoSPARC. **h**, Representative model segments fitted into EM density.

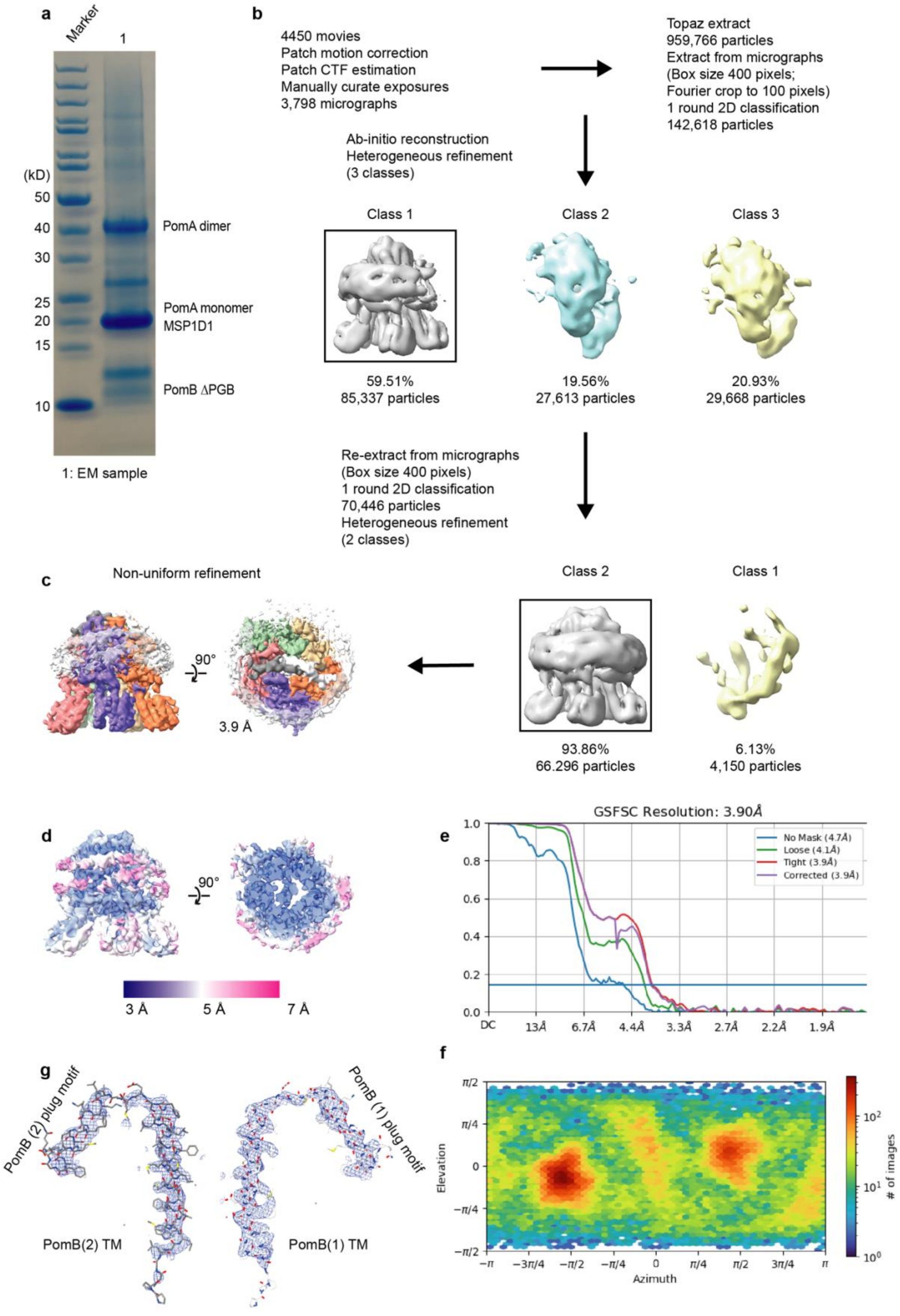
a
Marker
1
(kD)
50
40
30
25
20
15
10
PomA dimer
PomA monomer
MSP1D1
PomB ΔPGB
1: EM sample
b
4450 movies
Patch motion correction
Patch CTF estimation
Manually curate exposures
3,798 micrographs
Topaz extract
959,766 particles
Extract from micrographs
(Box size 400 pixels;
Fourier crop to 100 pixels)
1 round 2D classification
142,618 particles
Ab-initio reconstruction
Heterogeneous refinement
(3 classes)
Class 1
Class 2
Class 3
59.51%
85,337 particles
19.56%
27,613 particles
20.93%
29,668 particles
Re-extract from micrographs
(Box size 400 pixels)
1 round 2D classification
70,446 particles
Heterogeneous refinement
(2 classes)
c
Non-uniform refinement
90°
3.9 Å
Class 2
Class 1
93.86%
66.296 particles
6.13%
4,150 particles
d
90°
3 Å
5 Å
7 Å
e
GSFSC Resolution: 3.90Å
No Mask (4.7Å)
Loose (4.1Å)
Tight (3.9Å)
Corrected (3.9Å)
DC
13Å
6.7Å
4.4Å
3.3Å
2.7Å
2.2Å
1.9Å
f
Elevation
Azimuth
### of images
g
PomB (2) plug motif
PomB (1) plug motif
PomB(2) TM
PomB(1) TM

**Fig. S3 Cryo-EM of *Va*PomAB in MSP1D1 lipid nanodisc.**

**a**, SDS gel analysis of purified *Va*PomAB in MSP1D1 lipid nanodisc. **b**, Flowchart of the data processing of *Va*PomAB in MSP1D1 lipid nanodisc in cryoSPARC that results in the final cryo-EM structure. **c**, The final cryo-EM map of *Va*PomAB in MSP1D1 lipid nanodisc at around 3.9 Å resolution. **d**, Cryo-EM density map of *Va*PomAB in MSP1D1 lipid nanodisc colored by local resolution (in Å) estimated in cryoSPARC. **e**, Gold standard (0.143) Fourier shell correlation (GSFSC) curves for *Va*PomAB in MSP1D1 lipid nanodisc. **f**, Particle directional distribution of *Va*PomAB in MSP1D1nanodisc. **g**, Representative model segments fitted into EM density.

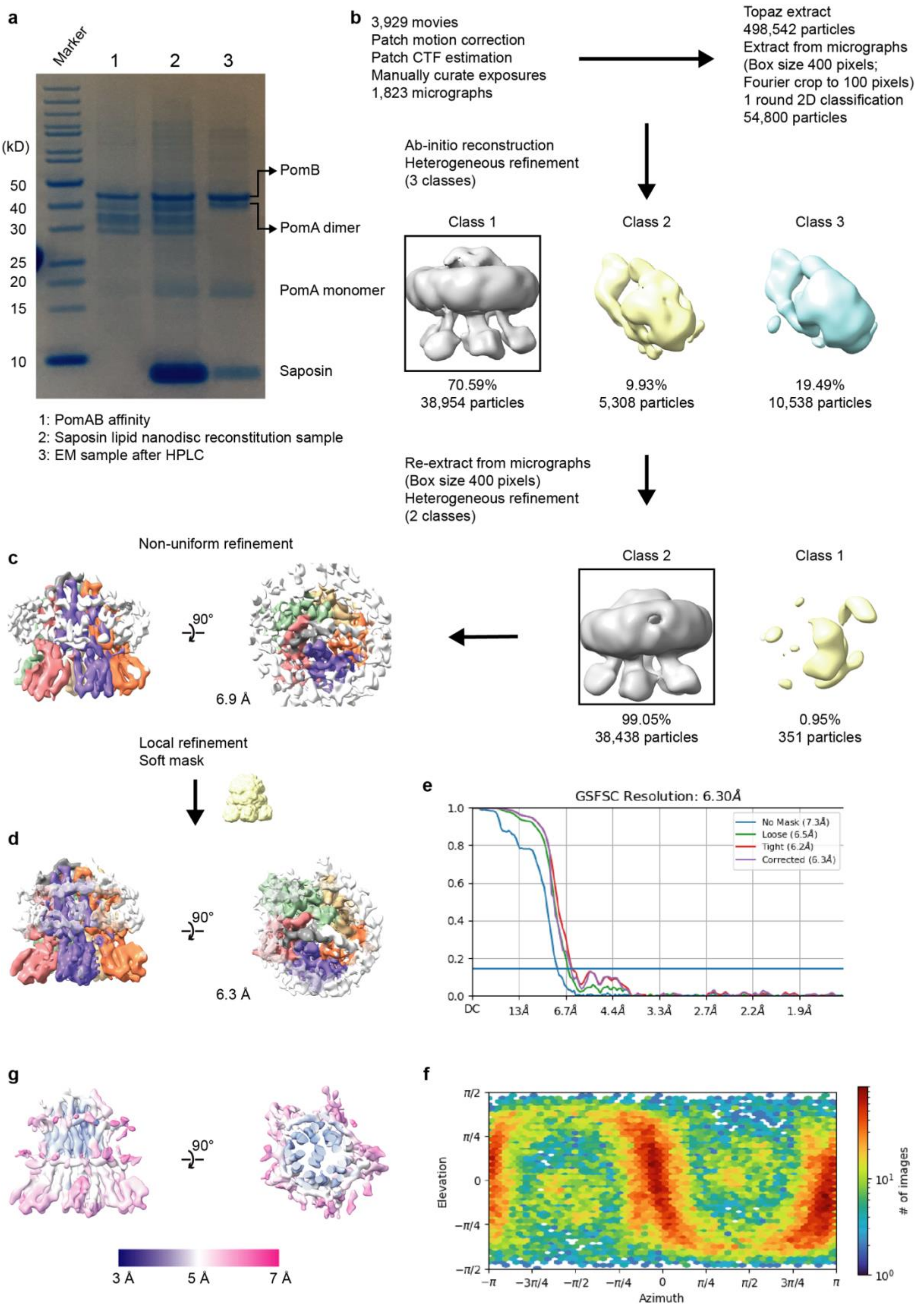
a
Marker
1
2
3
(kD)
50
40
30
25
20
15
10
PomB
PomA dimer
PomA monomer
Saposin
1: PomAB affinity
2: Saposin lipid nanodisc reconstitution sample
3: EM sample after HPLC
b
3,929 movies
Patch motion correction
Patch CTF estimation
Manually curate exposures
1,823 micrographs
Topaz extract
498,542 particles
Extract from micrographs
(Box size 400 pixels;
Fourier crop to 100 pixels)
1 round 2D classification
54,800 particles
Ab-initio reconstruction
Heterogeneous refinement
(3 classes)
Class 1
Class 2
Class 3
70.59%
38,954 particles
9.93%
5,308 particles
19.49%
10,538 particles
Re-extract from micrographs
(Box size 400 pixels)
Heterogeneous refinement
(2 classes)
Class 2
Class 1
99.05%
38,438 particles
0.95%
351 particles
c
Non-uniform refinement
90°
6.9 Å
Local refinement
Soft mask
d
90°
6.3 Å
e
GSFSC Resolution: 6.30Å
No Mask (7.3Å)
Loose (6.5Å)
Tight (6.2Å)
Corrected (6.3Å)
1.0
0.8
0.6
0.4
0.2
0.0
DC
13Å
6.7Å
4.4Å
3.3Å
2.7Å
2.2Å
1.9Å
g
90°
3 Å
5 Å
7 Å
f
π/2
π/4
0
−π/4
−π/2
Elevation
−π
−3π/4
−π/2
−π/4
0
π/4
π/2
3π/4
π
Azimuth
### of images
$10^1$
$10^0$

**Fig. S4 Cryo-EM of full length *Va*PomAB in saposin lipid nanodisc.**

**a**, SDS gel analysis of the purified full length *Va*PomAB in saposin lipid nanodisc. **b**-**c**, Flowchart of the data processing of full length *Va*PomAB in saposin lipid nanodisc in cryoSPARC that results in the final cryo-EM structure. **d**, The final cryo-EM map of *Va*PomAB in saposin lipid nanodisc at around 6.3 Å resolution after local refinement. **e**, Gold standard (0.143) Fourier shell correlation (GSFSC) curves for *Va*PomAB in saposin lipid nanodisc. **f**, Particle directional distribution. **g**, Cryo-EM density map of *Va*PomAB in saposin lipid nanodisc colored by local resolution (in Å) estimated in cryoSPARC.

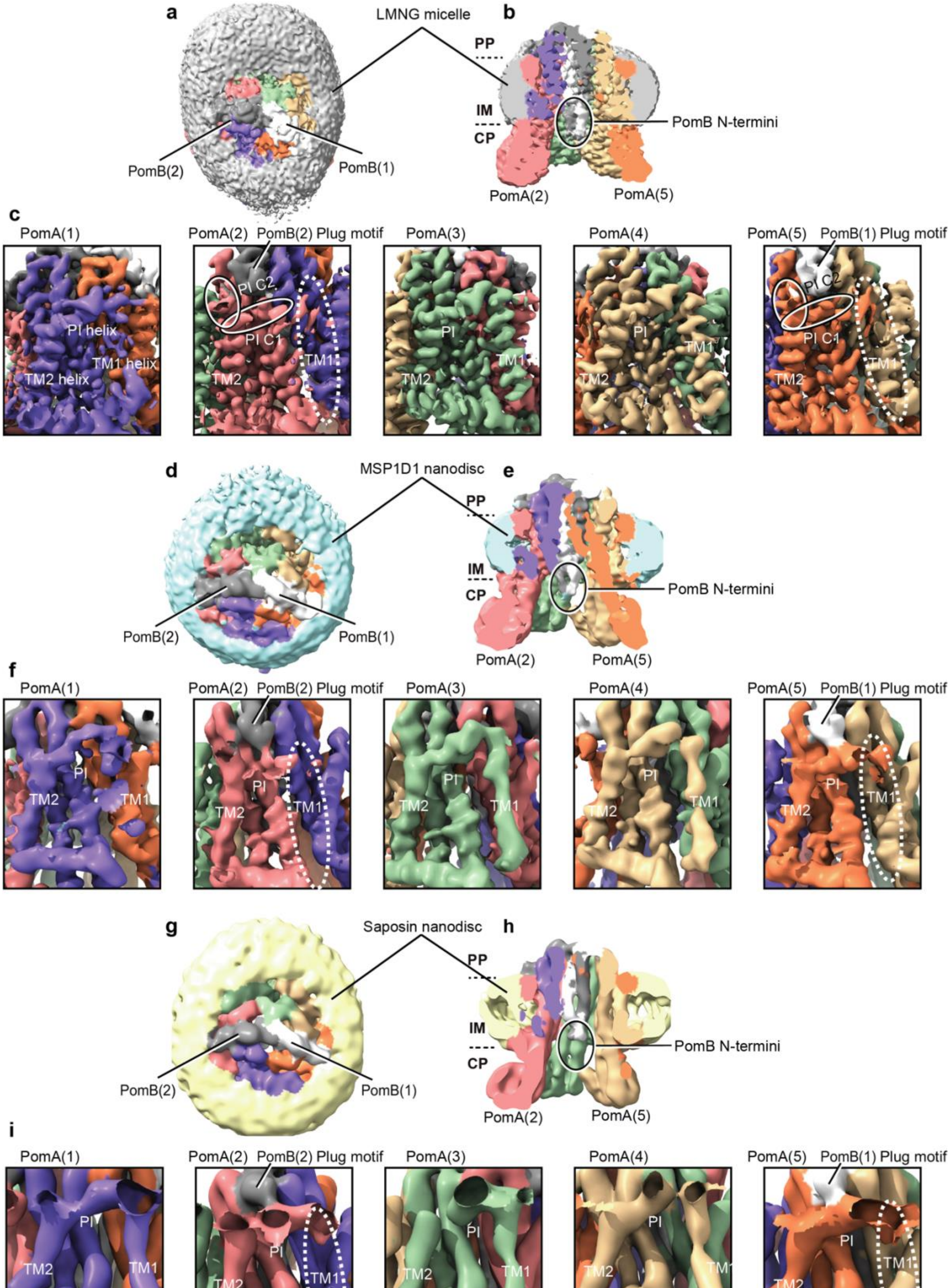
a
LMNG micelle
b
PP
IM
CP
PomB N-termini
PomB(2)
PomB(1)
PomA(2)
PomA(5)
c
PomA(1)
PomA(2)
PomB(2) Plug motif
PomA(3)
PomA(4)
PomA(5)
PomB(1) Plug motif
PI helix
TM1 helix
TM2 helix
PI C2
PI C1
TM1
TM2
PI
d
MSP1D1 nanodisc
e
PP
IM
CP
PomB N-termini
PomB(2)
PomB(1)
PomA(2)
PomA(5)
f
PomA(1)
PomA(2)
PomB(2) Plug motif
PomA(3)
PomA(4)
PomA(5)
PomB(1) Plug motif
PI
TM2
TM1
g
Saposin nanodisc
h
PP
IM
CP
PomB N-termini
PomB(2)
PomB(1)
PomA(2)
PomA(5)
i
PomA(1)
PomA(2)
PomB(2) Plug motif
PomA(3)
PomA(4)
PomA(5)
PomB(1) Plug motif
PI
TM2
TM1

**Fig. S5 Dynamics of *Va*PomA PI and TM1 helices.**

**a**-**c**, Representation of the *Va*PomAB LMNG unsharpened electrostatic potential maps at low threshold showing the conformational dynamic of PI helices that interact with PomB plug motifs, and the flexibility of the corresponding TM1 helices. **d**-**f**, Representation of the *Va*PomAB MSP1D1 lipid nanodisc unsharpened electrostatic potential maps at low threshold. **g**-**i**, Representation of the full length *Va*PomAB saposin lipid nanodisc unsharpened electrostatic potential maps at low threshold.

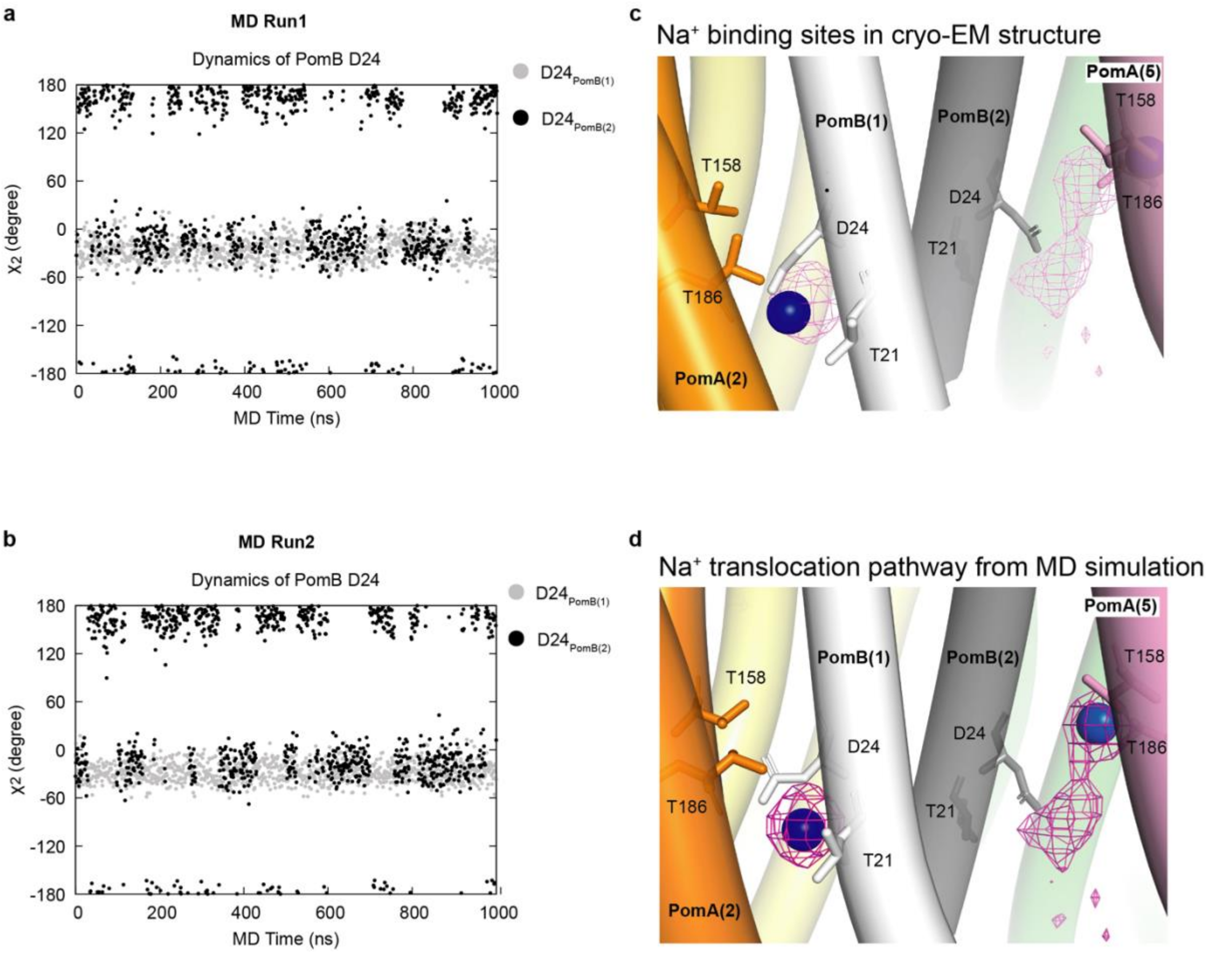


**Fig. S6 Na$^+$ translocation pathway and dynamics of PomB D24.**

**a**-**b**, The trajectories of the side chain dynamics of D24 in PomB chain 1 and 2 obtained from two independent MD simulations. **c**, The cryo-EM Na$^+$ binding sites. The modelled Na$^+$ ions are shown by blue spheres. **d**, The Na$^+$ binding sites captured in MD simulations. The average density of Na$^+$ ions is represented by red mesh in **c** and **d**.

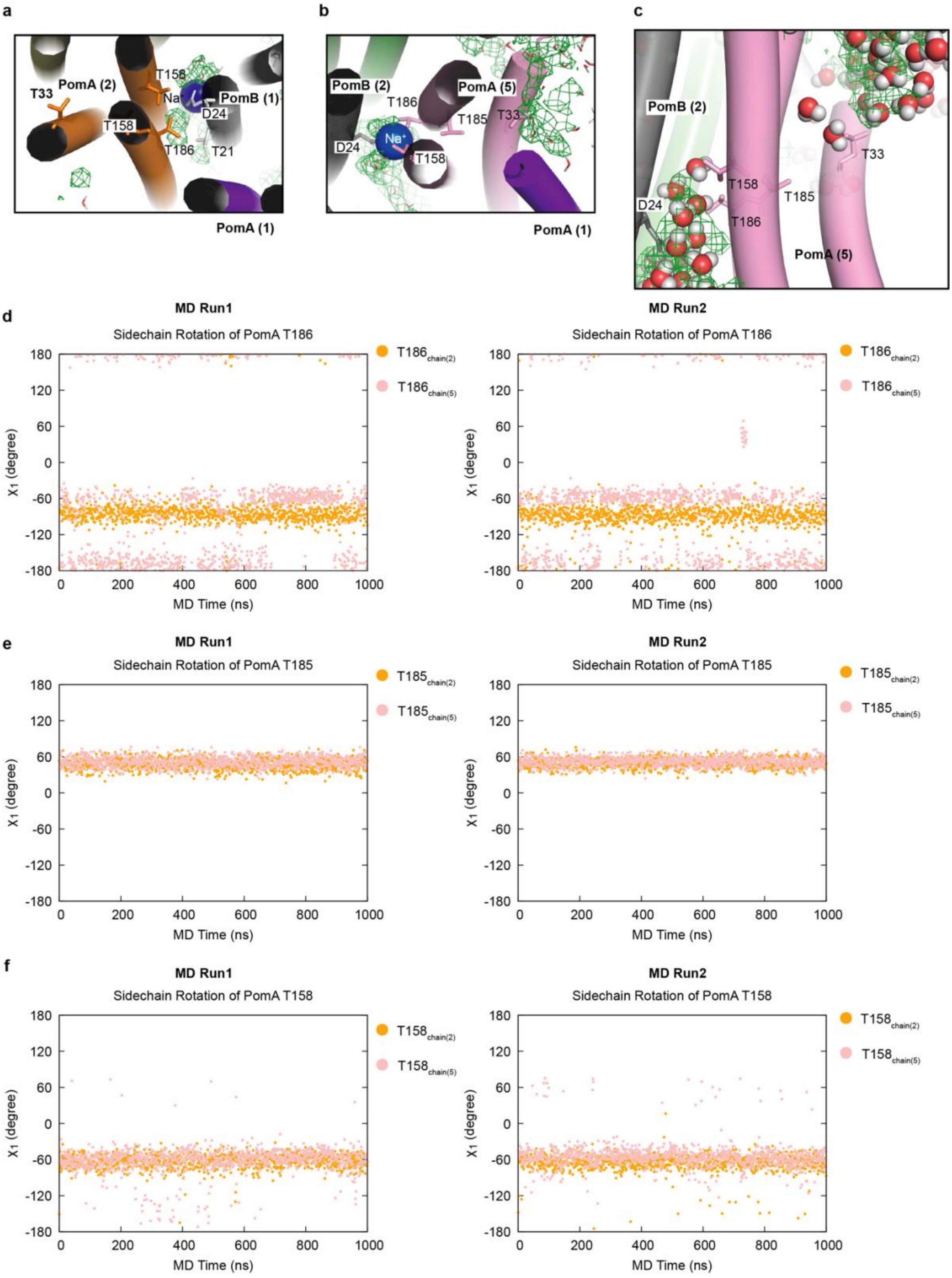

a
T33
PomA (2)
T158
Na
PomB (1)
D24
T158
T186
T21
PomA (1)
b
PomB (2)
T186
PomA (5)
T185
T33
D24
Na+
T158
PomA (1)
c
PomB (2)
T33
T158
T185
D24
T186
PomA (5)
d
MD Run1
Sidechain Rotation of PomA T186
T186 chain(2)
T186 chain(5)
X1 (degree)
MD Time (ns)
MD Run2
Sidechain Rotation of PomA T186
e
MD Run1
Sidechain Rotation of PomA T185
T185 chain(2)
T185 chain(5)
MD Run2
Sidechain Rotation of PomA T185
f
MD Run1
Sidechain Rotation of PomA T158
T158 chain(2)
T158 chain(5)
MD Run2
Sidechain Rotation of PomA T158

**Fig. S7 Hydration of T33 and the $Na^+$ translocation pathway and side chain dynamics of T158, T185 and T186 obtained from explicit solvent MD simulations.**

**a**-**b**, The hydration and $Na^+$ binding in the engaged and disengaged state, respectively. The average density of water molecules is represented by mesh in green. **c**, A snapshot from the MD simulations to show the hydration of T33 in PomA chain 5. **d**-**f**, The MD trajectories of the side chain dynamics of T186, T185 and T158 in PomA chain 2 and 5.

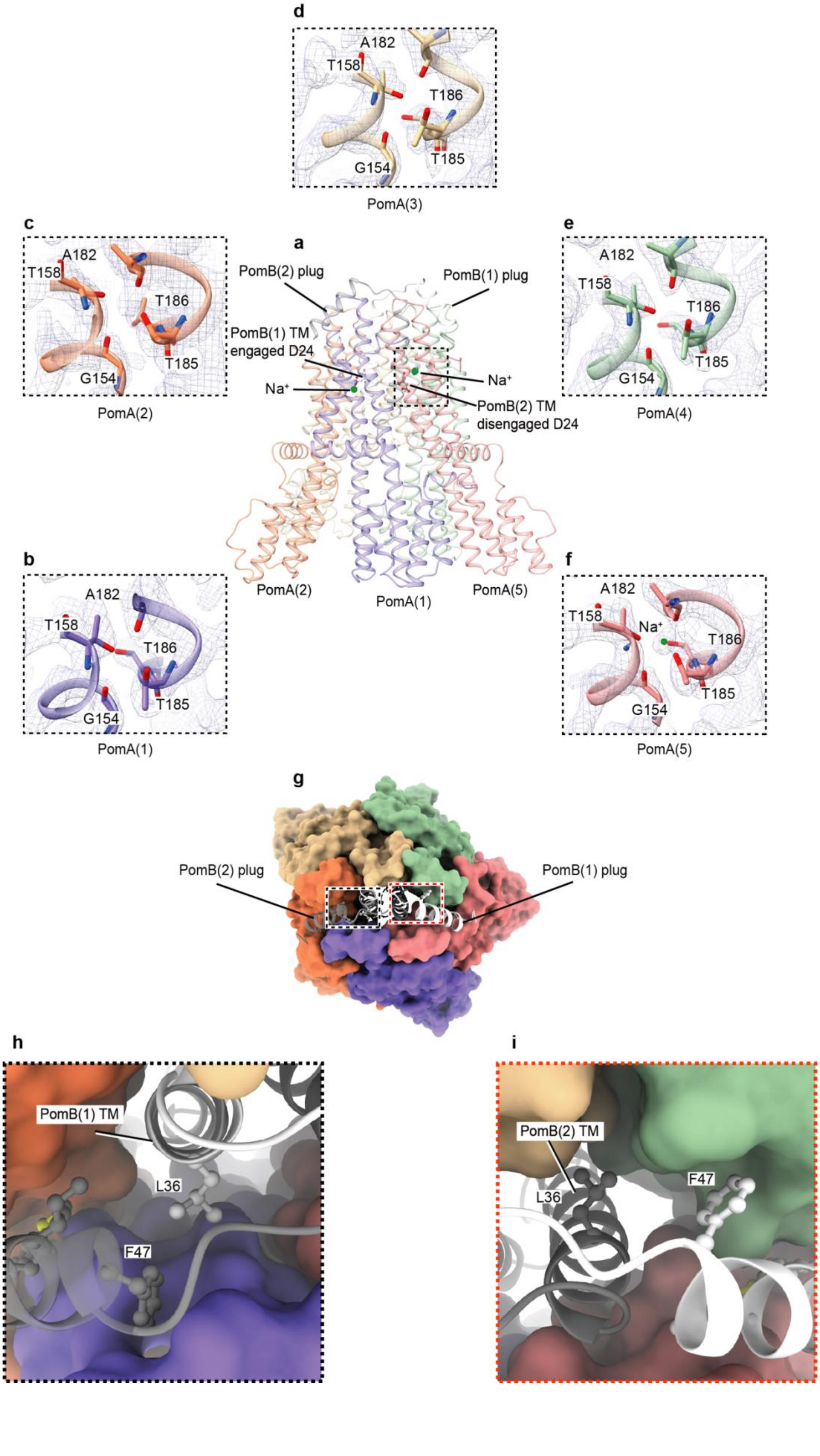

d
A182
T158
T186
T185
G154
PomA(3)
c
A182
T158
T186
T185
G154
PomA(2)
a
PomB(2) plug
PomB(1) plug
PomB(1) TM
engaged D24
Na⁺
Na⁺
PomB(2) TM
disengaged D24
PomA(2)
PomA(1)
PomA(5)
e
A182
T158
T186
T185
G154
PomA(4)
b
A182
T158
T186
T185
G154
PomA(1)
f
A182
T158
Na⁺
T186
T185
G154
PomA(5)
g
PomB(2) plug
PomB(1) plug
h
PomB(1) TM
L36
F47
i
PomB(2) TM
L36
F47

**Fig. S8 densities of ion selectivity cavities and PomB L36 interaction environments.**

**a**, View from the plane of the membrane, showing the position of ion selectivity cavity within the complex. **b-f**, ion selectivity cavities from PomA chains 1 to 5. EM densities are overlaid on the corresponding local regions. **g-i**, L36 from PomB chain 1 and chain 2 interaction environments, showing that PomB chain 1 L36 interacts PomB chain 2 F47.

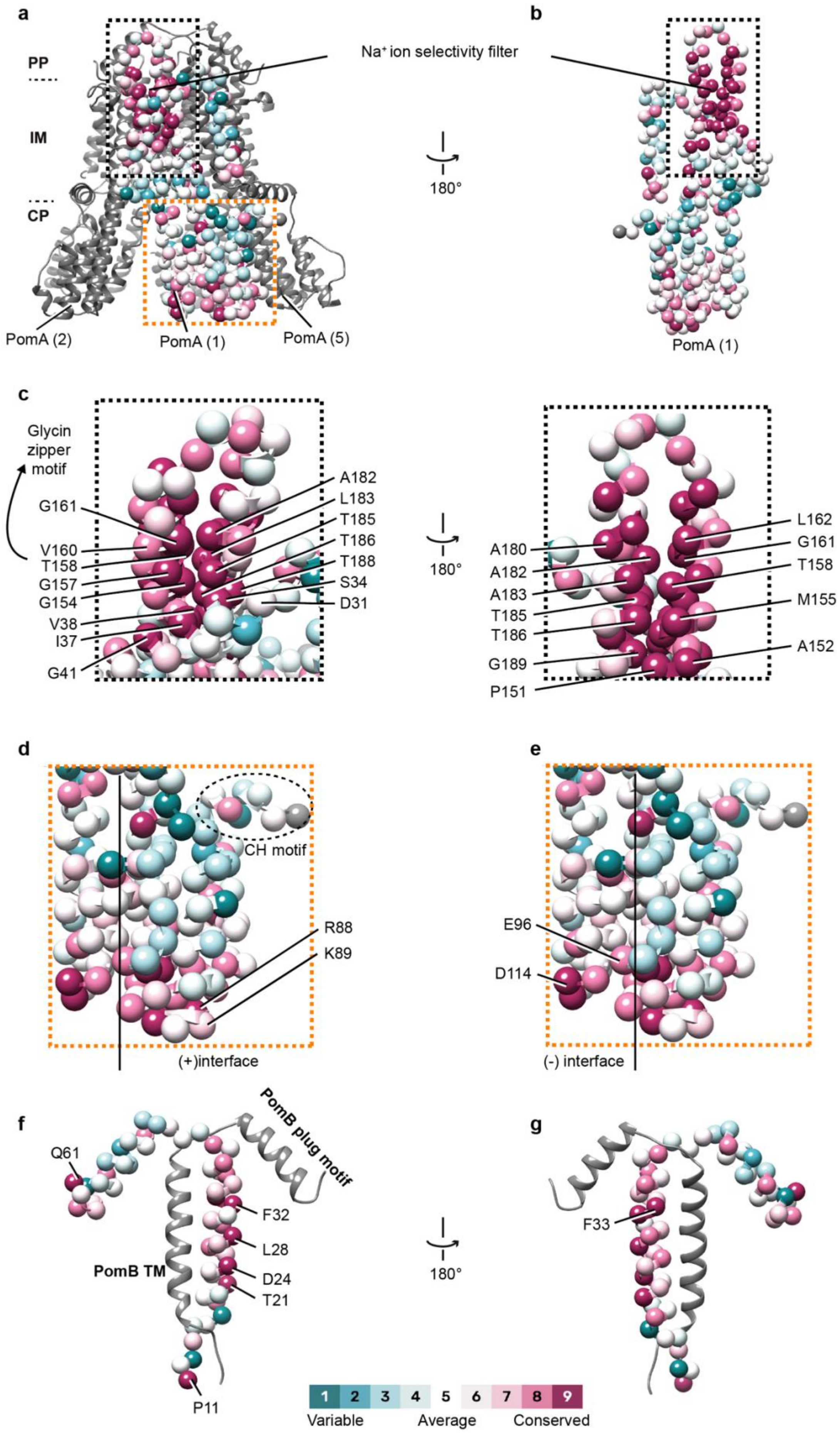
a
PP
IM
CP
PomA (2)
PomA (1)
PomA (5)
Na⁺ ion selectivity filter
180°
b
PomA (1)
c
Glycin zipper motif
G161
V160
T158
G157
G154
V38
I37
G41
A182
L183
T185
T186
T188
S34
D31
180°
A180
A182
A183
T185
T186
G189
P151
L162
G161
T158
M155
A152
d
CH motif
R88
K89
(+)interface
e
E96
D114
(-) interface
f
Q61
PomB plug motif
F32
L28
D24
T21
PomB TM
P11
180°
g
F33
1
2
3
4
5
6
7
8
9
Variable
Average
Conserved

**Fig. S9 Conservation (calculated with ConSurf) analysis of *Va*PomA and *Va*PomB.**

**a-b**, Conservation (calculated with ConSurf) of the surface residues of *Va*PomA from external and internal sides; Cα atom representation (shown as spheres) of the model colored by conservation. **c**, Conservation of the residues of the $Na^+$ ion selectivity filter and permeation pathway from the periplasmic side, both external and internal views are shown. **d**, Conservation of the residues of PomA cytoplasmic domain, highlighting the locations of the positively charged residues from the principal face involved in FliG torque helix binding. **e**, Same as in **d**, but highlighting negatively charged residues from the complementary face. **f**, Conservation of the surface residues of *Va*PomB, highlighting the strictly conserved residues. **g**, Same as in **f**, but rotated 180 degrees.

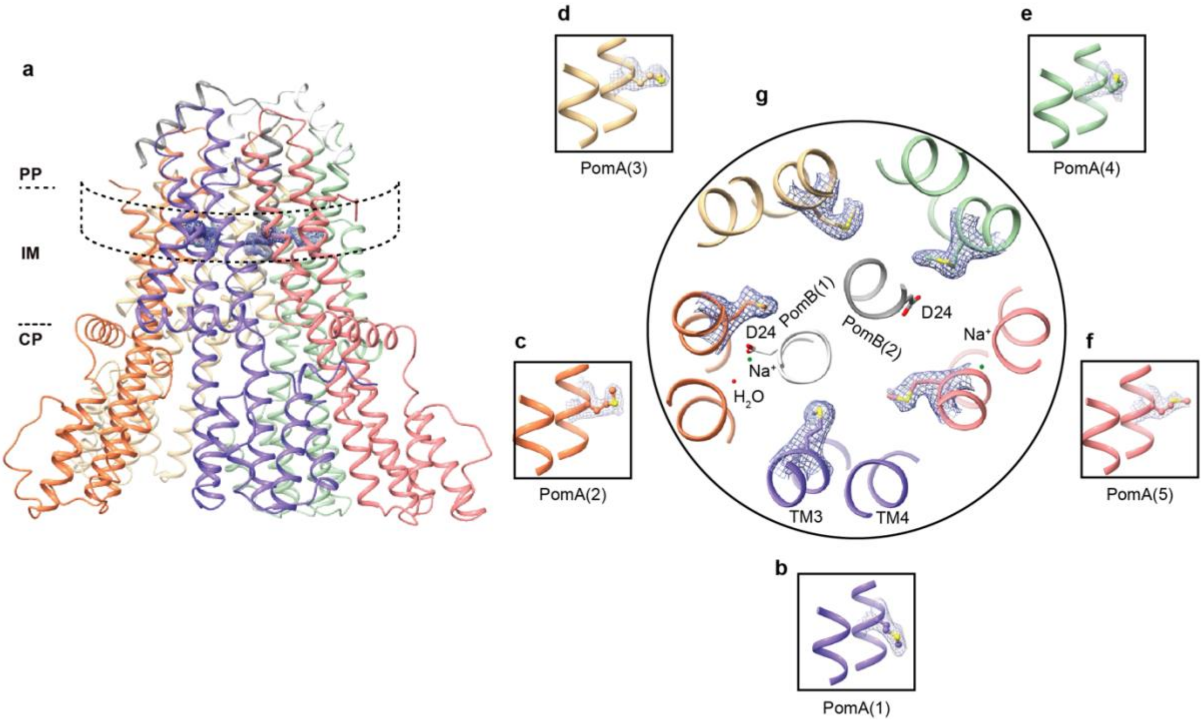


**Fig. S10 Conformational isomers of *Va*PomAB M155.**

**a**, View from the plane of the membrane, showing the position of PomA M155 within the complex. **b**-**f**, M155 isomers from PomA chains 1 to 5. EM densities are overlaid on the side chains of M155. **g**, Conformational isomers of M155 viewed from the top of the membrane.

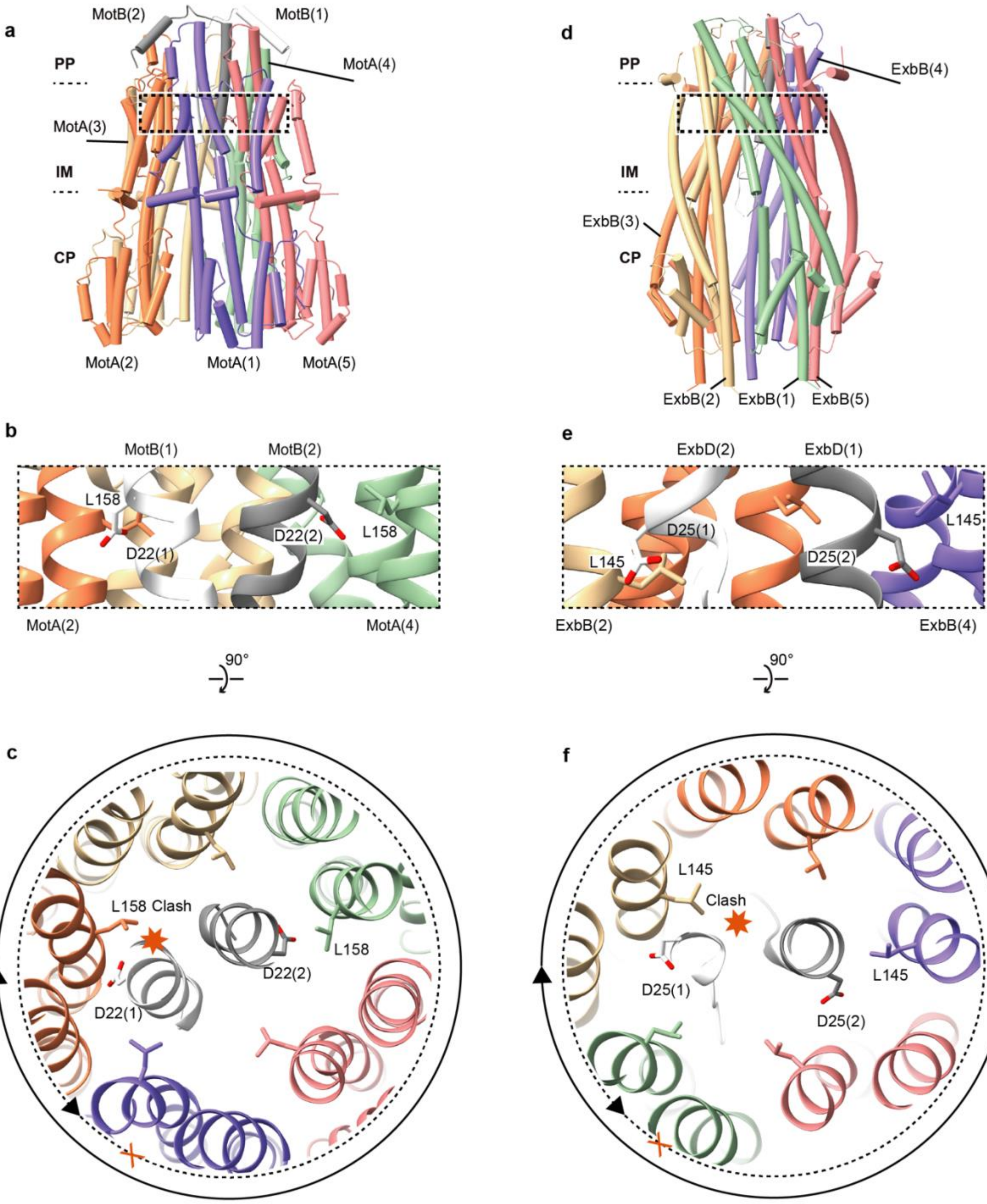


CjMotAB (PDB: 6YKM)

EcExbB-ExbD (PDB: 6TKI)

**Fig. S11 5:2 rotary motor directional rotation ‘reinforcement’ point.**

**a**, Proton-driven flagellar stator unit *Cj*MotAB (PDB: 6YKM). **b**, Conformational isomers of L158 near MotB engaged D24 and disengaged D24. **c**, Conformational isomers of L158 viewed from the top of the membrane. Solid circle indicates the rotational direction of MotA around MotB. The potential clash that would occur if PomA rotated CCW around PomB is indicated with a red heptagon. **d**, Proton-driven Ton ExbB-ExbD complex (PDB: 6TKI). **e**, Conformational isomers of L145 near ExbD engaged D25 and disengaged D25. **f**, Conformational isomers of ExbB L145 viewed from the top of the membrane. Solid circle indicates the rotational direction of ExbB around ExbD. The potential clash that would occur if ExbB rotated CCW around ExbD is indicated with a red heptagon.

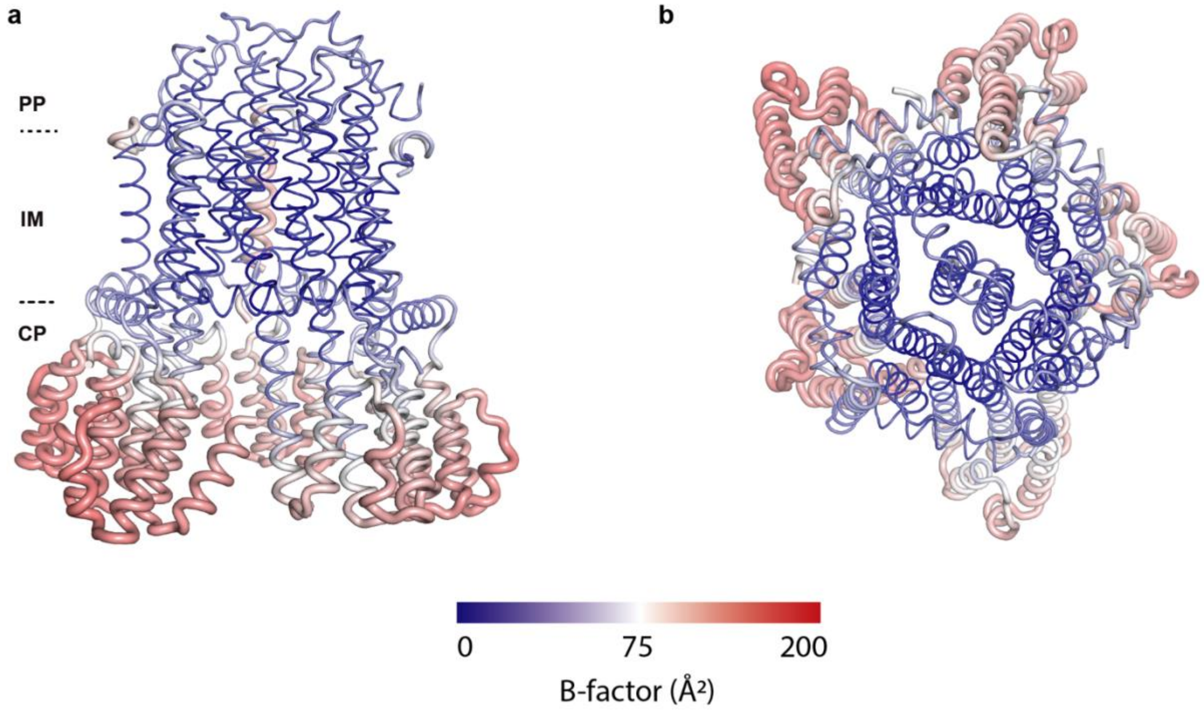


**Fig. S12 *Va*PomAB model B-factor distribution.**

Top (**a**) and side views (**b**) of the PomAB model (LMNG dataset) colored by B-factor distribution (atomic displacement factor).

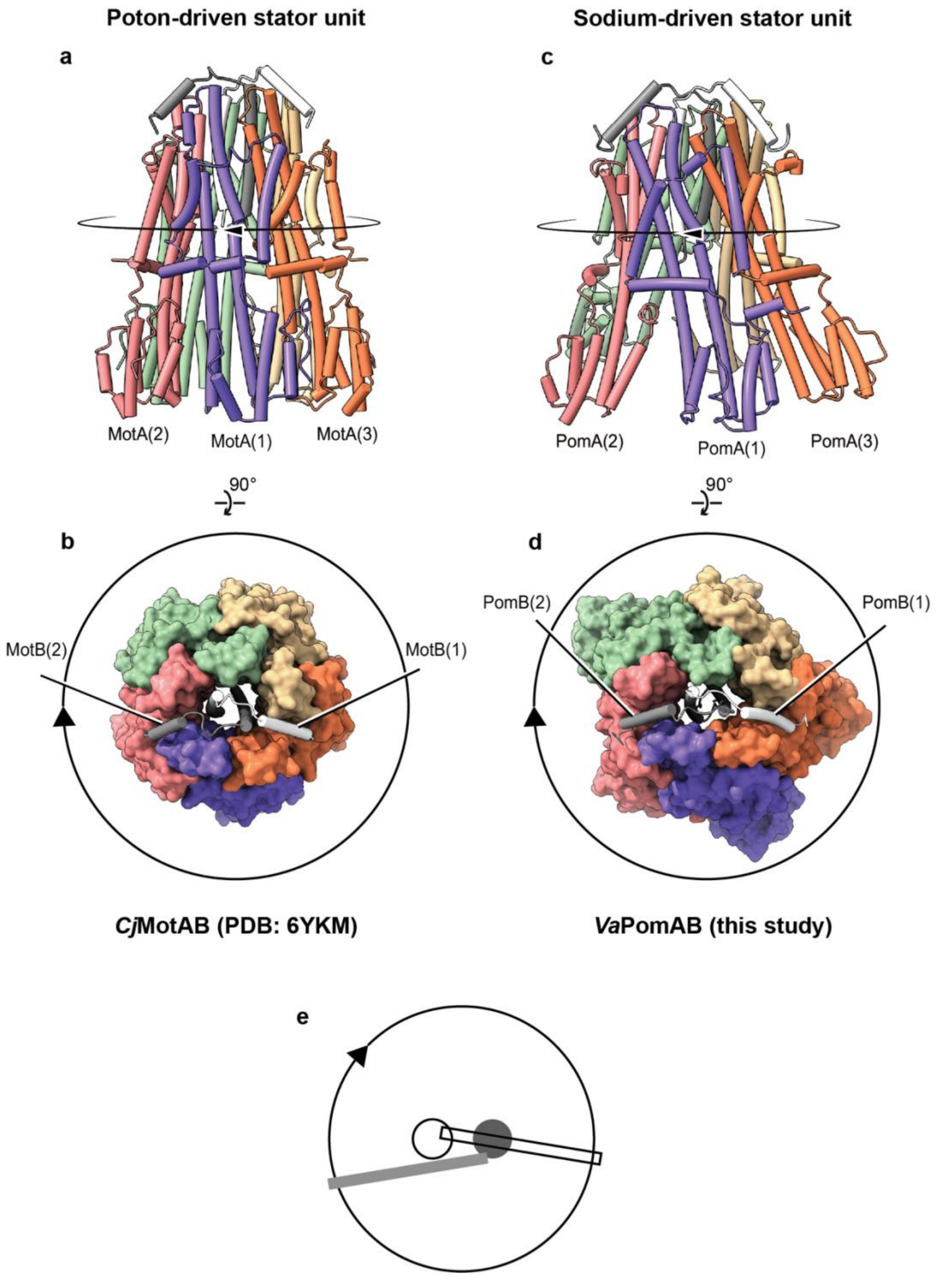


**Fig. S13 H⁺- and Na⁺- driven stator units PomB/MotB plug motifs organization.**

**a**, Side view of the proton-driven stator unit *Cj*MotAB in its auto-inhibited state.**b**, *Cj*MotAB viewed from the top of the membrane. **c**, Side view of the sodium-driven stator *Va*PomAB in its auto-inhibited state. **d**, *Va*PomAB viewed from the top of the membrane. Rotational direction of the stator unit is indicated. **e**, The unique trans mode organization of the plug motifs tightly blocks the CW rotation of the stator unit.

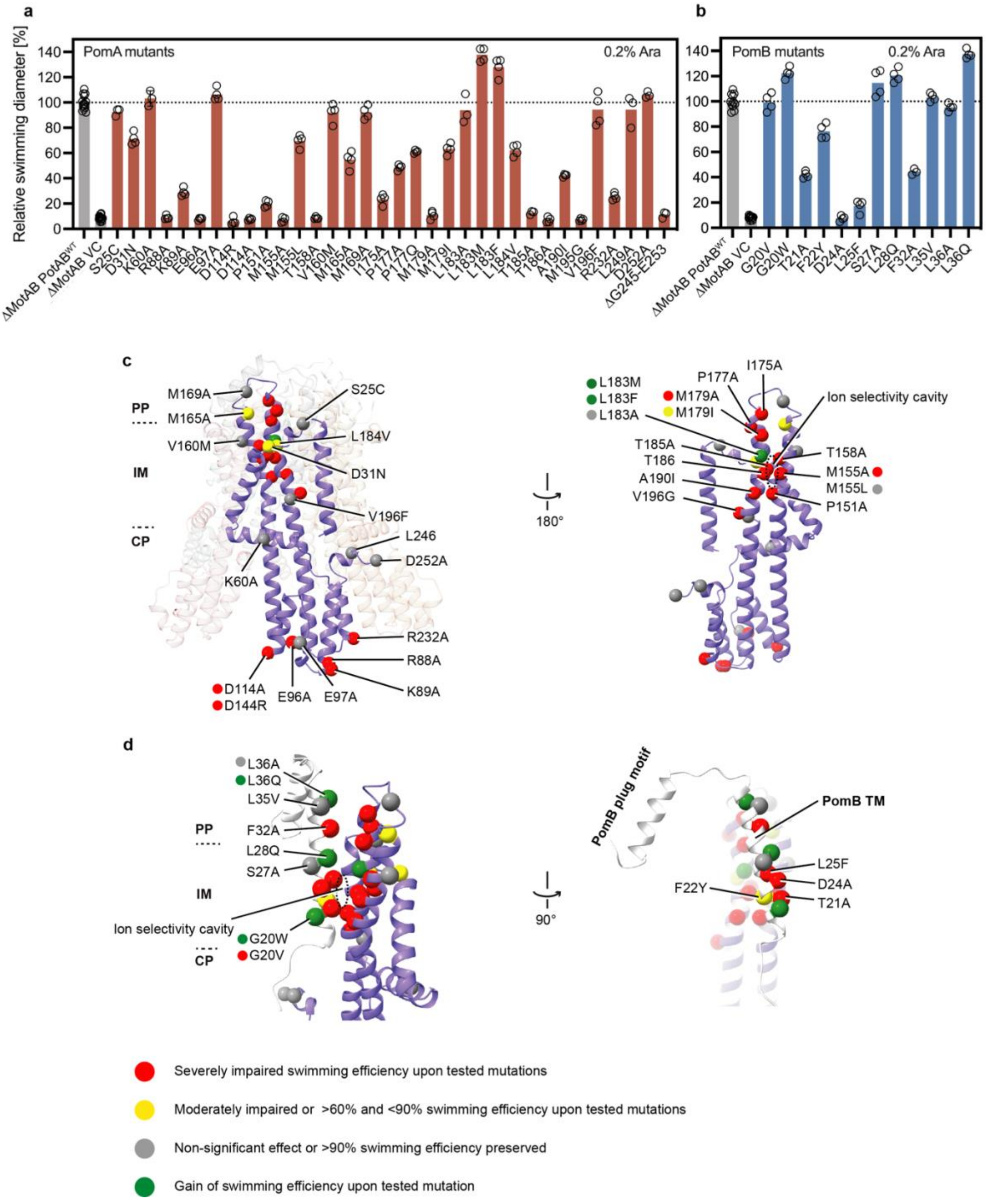


**Fig. S14 Mutational analysis for *Va*PomA and *Va*PomB plotted onto the *Va*PomAB structure.**

**a-b**, The motility phenotypes of VaPotAB PomA (**a**) and PotB (**b**) point mutants were analyzed using soft-agar motility plates containing 0.2% agar. **c-d**, Swimming efficiency of the *Va*PotAB point mutants, showing the mutated residues as Cα spheres on the PomA (purple) and PomB (white) structure.

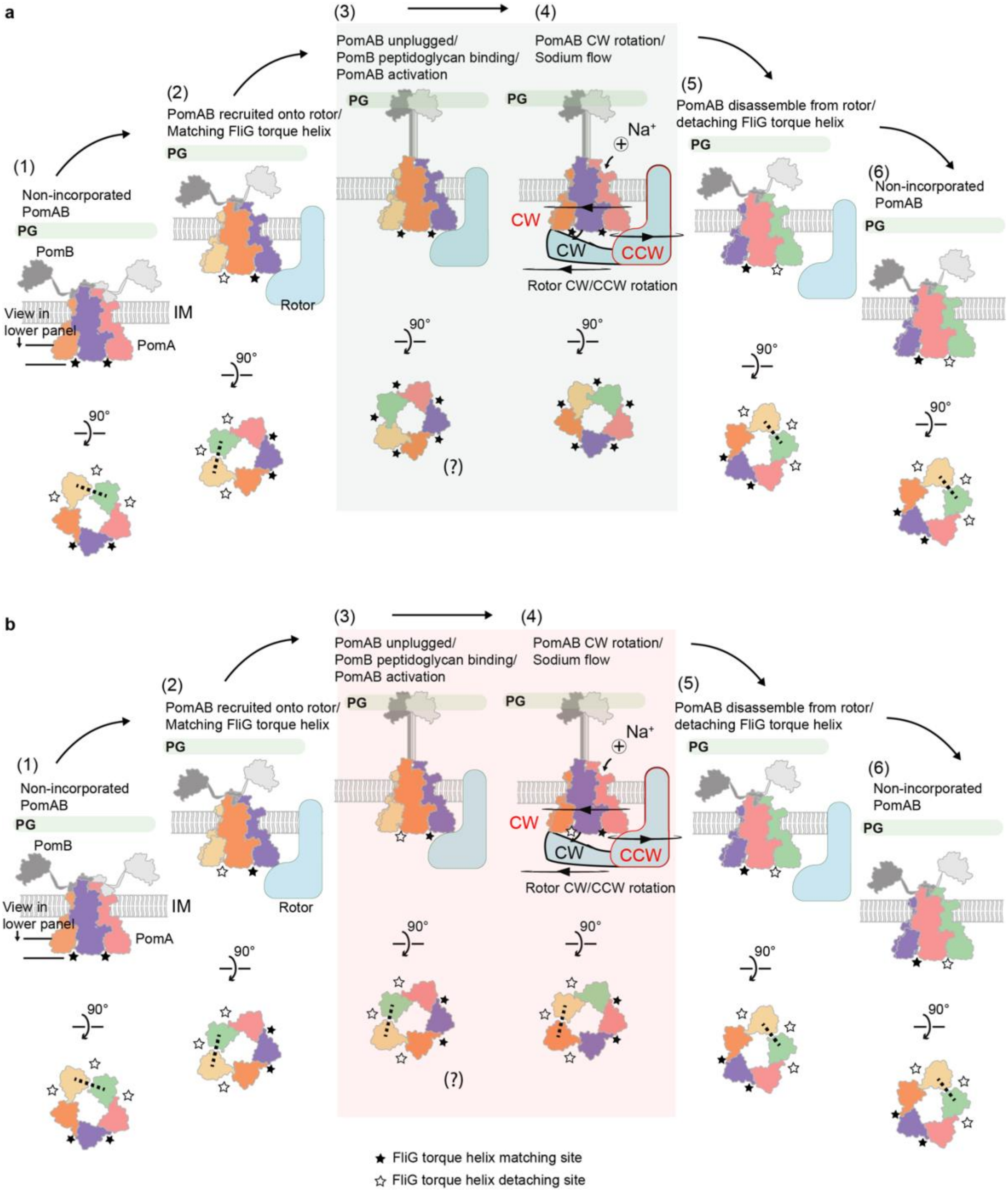
a
(1)
Non-incorporated PomAB
PG
PomB
IM
View in lower panel
PomA
90°
(2)
PomAB recruited onto rotor/ Matching FliG torque helix
PG
Rotor
90°
(3)
PomAB unplugged/ PomB peptidoglycan binding/ PomAB activation
PG
90°
(?)
(4)
PomAB CW rotation/ Sodium flow
PG
Na+
CW
CW
CCW
Rotor CW/CCW rotation
90°
(5)
PomAB disassemble from rotor/ detaching FliG torque helix
PG
90°
(6)
Non-incorporated PomAB
PG
90°
b
(1)
Non-incorporated PomAB
PG
PomB
IM
View in lower panel
PomA
90°
(2)
PomAB recruited onto rotor/ Matching FliG torque helix
PG
Rotor
90°
(3)
PomAB unplugged/ PomB peptidoglycan binding/ PomAB activation
PG
90°
(?)
(4)
PomAB CW rotation/ Sodium flow
PG
Na+
CW
CW
CCW
Rotor CW/CCW rotation
90°
(5)
PomAB disassemble from rotor/ detaching FliG torque helix
PG
90°
(6)
Non-incorporated PomAB
PG
90°
★ FliG torque helix matching site
☆ FliG torque helix detaching site

**Fig. S15 Models of conformational changes of PomA cytoplasmic domain during stator unit activation and disassembly from the rotor.**

**a**, PomA cytoplasmic domain is asymmetric, and one site of the CH-CI detachment is indicated in dashed line. Inactive stator unit orients its cytoplasmic domain towards the rotor to contact FliG torque helix through FliG torque helix ‘matching sites’ ((1)-(2)). During the activation, all five CH-CI interactions established, and PomA cytoplasmic domain becomes symmetric ((3)-(4)). The rotor could rotate either CW or CCW direction, depending on how it interacts with the stator unit. Stator unit disassembly from the rotor when external torque is decreased ((5)-(6)). **b**, In this model, during the stator unit activation, PomA cytoplasmic domain remains asymmetric ((3)-(4)); one site of the CI helix attaches to the PI helix and the adjacent CI helix detaches from the PI helix, sequentially creating a FliG torque helix ‘catching’ site that interacts with the FliG torque helix.
