## Supplemental tables for "Mechanisms of ion selectivity and rotor coupling in the bacterial flagellar sodium-driven stator unit"

### Supplemental Table S1: Cryo-EM data collection, refinement and validation statistics.

| | *Va*PomAB | *Va*PomAB (MSP1D1 nanodisc) | *Va*PomAB (Saposin nanodisc) |
|---|---|---|---|
| **Data collection and processing** | | | |
| Microscope | Titan Krios G2 | | |
| Voltage (kV) | 300 | | |
| Magnification (nominal) | 96,000x | | |
| Total exposure ($e^-/Å^2$) | 37.98 | 40.00 | 37.98 |
| Exposure fractions (no.) | 40 | | |
| Pixel size (Å) | 0.832 | | |
| Movies used (no.) | 6,444 | 3,798 | 1,823 |
| Total picked particles (no.) | 3,659,926 | 959,766 | 498,542 |
| Final particles (no.) | 923,963 | 66,296 | 38,438 |
| Box size (pixels) | 500 | 400 | 400 |
| Symmetry imposed | C1 | | |
| Map resolution (Å) (FSC 0.143) | 2.48 | 3.90 | 6.30 |
| **Refinement** | | | |
| Model composition | | | |
| Non-hydrogen atoms | 9,926 | 99,16 | |
| Protein residues | 1,312 | 1,312 | |
| Solvent molecules | 12 | - | |
| B-factors (mean; $Å^2$) | | | |
| Protein | 75.39 | 123.52 | |
| Solvent | 30.06 | - | |
| R.m.s. deviations | | | |
| Bond lengths (Å) | 0.006 | 0.006 | |
| Bond angles (□) | 0.874 | 1.288 | |
| CC (mask) | 0.83 | 0.79 | |
| Refinement resolution (FSC map vs. model (masked)=0.143) (Å) | 2.4 | 3.8 | |
| **Validation** | | | |
| MolProbity score | 2.04 | 2.08 | |
| Poor rotamers (%) | 0.00 | 0.09 | |
| Ramachandran plot | | | |
| Favored (%) | 97.61 | 97.23 | |
| Allowed (%) | 2.39 | 2.77 | |
| Disallowed (%) | 0.00 | 0.00 | |

### Supplemental Table S2: bacterial strains

| Number | Genotype | Source |
|---|---|---|
| n/a | NEB Dh5alpha (cloning strain) | NEB, Ipswich, MA, USA |
| TH437 | Salmonella enterica serovar Typhimurium LT2 | J. Roth (University of California, Davis) |
| EM10372 | DmotAB (leaving first and last 15 bp) | Mònica Santiveri et al. 2020 |
| EM12719 | LT2 / hhd-pbad33-pomapotb59-e-coli (p46) (CmR) | This study |
| EM12720 | LT2 / hhd-p47-pbad33-pomapotb59-samonella (p47) (CmR) | This study |
| EM12721 | DmotAB / hhd-pbad33-pomapotb59-e-coli (p46) (CmR) | This study |
| EM12722 | DmotAB / hhd-p47-pbad33-pomapotb59-samonella (p47) (CmR) | This study |
| EM12734 | LT2 / pBAD33.1 (Adgene #36267; CmR) | This study |
| EM12735 | DmotAB / pBAD33.1 (Adgene #36267; CmR) | This study |
| EM13301 | DmotAB / pEM13101 (pBAD33.1-pomA(S25C)pomB, Samonella PG-binding domain, CmR) | This study |
| EM13302 | DmotAB / pEM13102 (pBAD33.1-pomA(D31N)pomB, Samonella PG-binding domain, CmR) | This study |
| EM13303 | DmotAB / pEM13103 (pBAD33.1-pomA(R88A)pomB, Samonella PG-binding domain, CmR) | This study |
| EM13304 | DmotAB / pEM13104 (pBAD33.1-pomA(K89A)pomB, Samonella PG-binding domain, CmR) | This study |
| EM13305 | DmotAB / pEM13105 (pBAD33.1-pomA(E96A)pomB, Samonella PG-binding domain, CmR) | This study |
| EM13306 | DmotAB / pEM13106 (pBAD33.1-pomA(E97A)pomB, Samonella PG-binding domain, CmR) | This study |
| EM13307 | DmotAB / pEM13107 (pBAD33.1-pomA(D114R)pomB, Samonella PG-binding domain, CmR) | This study |
| EM13308 | DmotAB / pEM13108 (pBAD33.1-pomA(D114A)pomB, Samonella PG-binding domain CmR) | This study |
| EM13309 | DmotAB / pEM13109 (pBAD33.1-pomA(P151A)pomB, Samonella PG-binding domain, CmR) | This study |
| EM13310 | DmotAB / pEM13110 (pBAD33.1-pomA(M155A)pomB, Samonella PG-binding domain, CmR) | This study |
| EM13311 | DmotAB / pEM13111 (pBAD33.1-pomA(M155L)pomB, Samonella PG-binding domain, CmR) | This study |
| EM13312 | DmotAB / pEM13112 (pBAD33.1-pomA(T158A)pomB, Samonella PG-binding domain, CmR) | This study |
| EM13313 | DmotAB / pEM13113 (pBAD33.1-pomA(V160M)pomB, Samonella PG-binding domain, CmR) | This study |
| EM13314 | DmotAB / pEM13114 (pBAD33.1-pomA(M165A)pomB, Samonella PG-binding domain, CmR) | This study |
| EM13315 | DmotAB / pEM13115 (pBAD33.1-pomA(M169A)pomB, Samonella PG-binding domain, CmR) | This study |
| EM13316 | DmotAB / pEM13116 (pBAD33.1-pomA(I175A)pomB, Samonella PG-binding domain, CmR) | This study |
| EM13317 | DmotAB / pEM13117 (pBAD33.1-pomA(P177A)pomB, Samonella PG-binding domain, CmR) | This study |
| EM13318 | DmotAB / pEM13118 (pBAD33.1-pomA(P177Q)pomB, Samonella PG-binding domain, CmR) | This study |
| EM13319 | DmotAB / pEM13119 (pBAD33.1-pomA(M179A)pomB, Samonella PG-binding domain, CmR) | This study |
| EM13320 | DmotAB / pEM13120 (pBAD33.1-pomA(M179I)pomB, Samonella PG-binding domain, CmR) | This study |
| EM13321 | DmotAB / pEM13121 (pBAD33.1-pomA(L183A)pomB, Samonella PG-binding domain, CmR) | This study |
| EM13322 | DmotAB / pEM13122 (pBAD33.1-pomA(L183M)pomB, Samonella PG-binding domain, CmR) | This study |
| EM13323 | DmotAB / pEM13123 (pBAD33.1-pomA(L183F)pomB, Samonella PG-binding domain, CmR) | This study |
| EM13324 | DmotAB / pEM13124 (pBAD33.1-pomA(L184V)pomB, Samonella PG-binding domain, CmR) | This study |
| EM13325 | DmotAB / pEM13125 (pBAD33.1-pomA(T186A)pomB, Samonella PG-binding domain, CmR) | This study |
| EM13326 | DmotAB / pEM13126 (pBAD33.1-pomA(A190I)pomB, Samonella PG-binding domain, CmR) | This study |
| EM13327 | DmotAB / pEM13127 (pBAD33.1-pomA(M195G)pomB, Samonella PG-binding domain, CmR) | This study |
| EM13328 | DmotAB / pEM13128 (pBAD33.1-pomA(V196F)pomB, Samonella PG-binding domain, CmR) | This study |
| EM13329 | DmotAB / pEM13129 (pBAD33.1-pomA(R232A)pomB, Samonella PG-binding domain, CmR) | This study |
| EM13330 | DmotAB / pEM13130 (pBAD33.1-pomApomB(G20V), Samonella PG-binding domain, CmR) | This study |
| EM13331 | DmotAB / pEM13131 (pBAD33.1-pomApomB(G20W), Samonella PG-binding domain, CmR) | This study |
| EM13332 | DmotAB / pEM13132 (pBAD33.1-pomApomB(T21A), Samonella PG-binding domain, CmR) | This study |
| EM13333 | DmotAB / pEM13133 (pBAD33.1-pomApomB(F22Y), Samonella PG-binding domain, CmR) | This study |

| | | |
|---|---|---|
| EM13334 | DmotAB / pEM13134 (pBAD33.1-pomApomB(D24A), Samonella PG-binding domain, CmR) | This study |
| EM13335 | DmotAB / pEM13135 (pBAD33.1-pomApomB(L25F), Samonella PG-binding domain, CmR) | This study |
| EM13336 | DmotAB / pEM13136 (pBAD33.1-pomApomB(S27A), Samonella PG-binding domain, CmR) | This study |
| EM13337 | DmotAB / pEM13137 (pBAD33.1-pomApomB(L28Q), Samonella PG-binding domain, CmR) | This study |
| EM13338 | DmotAB / pEM13138 (pBAD33.1-pomApomB(F32A), Samonella PG-binding domain, CmR) | This study |
| EM13339 | DmotAB / pEM13139 (pBAD33.1-pomApomB(L35V), Samonella PG-binding domain, CmR) | This study |
| EM13340 | DmotAB / pEM13140 (pBAD33.1-pomApomB(L36A), Samonella PG-binding domain, CmR) | This study |
| EM13341 | DmotAB / pEM13141 (pBAD33.1-pomApomB(L36Q), Samonella PG-binding domain, CmR) | This study |
| EM13783 | DmotAB / pEM13778 (pBAD33.1-pomA(K60A)pomB, Samonella PG-binding domain CmR) | This study |
| EM13784 | DmotAB / pEM13779 (pBAD33.1-pomA(T185A)pomB, Samonella PG-binding domain CmR) | This study |
| EM13785 | DmotAB / pEM13780 (pBAD33.1-pomA(L249A)pomB, Samonella PG-binding domain CmR) | This study |
| EM13786 | DmotAB / pEM13781 (pBAD33.1-pomA(D252A)pomB, Samonella PG-binding domain CmR) | This study |
| EM13787 | DmotAB / pEM13782 (pBAD33.1-pomA(ΔAA245-253)pomB, Samonella PG-binding domain CmR) | This study |

### Supplemental Table S3: Primers

| Number | Name | Sequence 5'-3' |
|---|---|---|
| 5593 | Va-PomAS25C-fwd | GCTTGGTGGCtGCATCGGCATGTTTGTC |
| 5594 | Va-PomAS25C-rv | GACAAACATGCCGATGCaGCCACCAAGC |
| 5595 | Va-PomAD31N-fwd | CATGTTTGTCaATGTCACGTCGATCC |
| 5596 | Va-PomAD31N-rv | GGATCGACGTGACATtGACAAACATG |
| 5597 | Va-PomAR88A-fwd | GATGCGGCGgccAAAGGTGGTTTTCTTG |
| 5598 | Va-PomAR88A-rv | CAAGAAAACCACCTTTggcCGCCGCATC |
| 5599 | Va-PomAK89A-fwd | GATGCGGCGCGTgcgGGTGGTTTTC |
| 5600 | Va-PomAK89A-rv | GAAAACCACCcgcACGCGCCGCATC |
| 5601 | Va-PomAE96A-fwd | CTTGCTCTTGcgGAGATGGAAATAAAC |
| 5602 | Va-PomAE96A-rv | GTTTATTTCCATCTCcgCAAGAGCAAG |
| 5603 | Va-PomAE97A-fwd | CTTGCTCTTGAAGccATGGAAATAAAC |
| 5604 | Va-PomAE97A-rv | GTTTATTTCCATggCTTCAAGAGCAAG |
| 5605 | Va-PomAD114R-fwd | GATCTACTGGTTcgcGGCCATGATGC |
| 5606 | Va-PomAD114R-rv | GCATCATGGCCgcgAACCAGTAGATC |
| 5607 | Va-PomAD114A-fwd | GATCTACTGGTTGcgGGCCATGATG |
| 5608 | Va-PomAD114A-rv | CATCATGGCCcgCAACCAGTAGATC |
| 5609 | Va-PomAP151A-fwd | GGCGACGTTGCTgCcGCGATGGGAATG |
| 5610 | Va-PomAP151A-rv | CATTCCCATCGCgGcAGCAACGTCGCC |
| 5611 | Va-PomAM155A-fwd | CTGCGATGGGAgcGATTGGCACCTTG |
| 5612 | Va-PomAM155A-rv | CAAGGTGCCAATCgcTCCCATCGCAG |
| 5613 | Va-PomAM155L-fwd | CTGCGATGGGAcTGATTGGCACCTTG |
| 5614 | Va-PomAM155L-rv | CAAGGTGCCAATCAgTCCCATCGCAG |
| 5615 | Va-PomAT158A-fwd | GGGAATGATTGGCgCgTTGGTTGGTC |
| 5616 | Va-PomAT158A-rv | GACCAACCAAcGcGCCAATCATTCCC |
| 5617 | Va-PomAV160M-fwd | GATTGGCACCTTGaTgGGTCTTGTTG |
| 5618 | Va-PomAV160M-rv | CAACAAGACCcAtCAAGGTGCCAATC |
| 5619 | Va-PomAM165A-fwd | GGTCTTGTTGCGgcGCTTTCAAACATG |
| 5620 | Va-PomAM165A-rv | CATGTTTGAAAGCgcCGCAACAAGACC |
| 5621 | Va-PomAM169A-fwd | GCTTTCAAACgcGGATGACCCTAAAGC |
| 5622 | Va-PomAM169A-rv | GCTTTAGGGTCATCCgcGTTTGAAAGC |
| 5623 | Va-PomAI175A-fwd | GGATGACCCTAAAGCGgcgGGACCAGC |
| 5624 | Va-PomAI175A-rv | GCTGGTCCcgcCGCTTTAGGGTCATCC |
| 5625 | Va-PomAP177A-fwd | CTAAAGCGATTGGAgCAGCAATGGCCG |
| 5626 | Va-PomAP177A-rv | CGGCCATTGCTGcTCCAATCGCTTTAG |
| 5627 | Va-PomAP177Q-fwd | CTAAAGCGATTGGACaAGCAATGGCCG |
| 5628 | Va-PomAP177Q-rv | CGGCCATTGCTtGTCCAATCGCTTTAG |
| 5629 | Va-PomAM179A-fwd | GATTGGACCAGCAgcGGCCGTTGCAC |
| 5630 | Va-PomAM179A-rv | GTGCAACGGCCgcTGCTGGTCCAATC |
| 5631 | Va-PomAM179I-fwd | GATTGGACCAGCAATtGCCGTTGCAC |
| 5632 | Va-PomAM179I-rv | GTGCAACGGCaATTGCTGGTCCAATC |
| 5633 | Va-PomAL183A-fwd | GGCCGTTGCAgcCTTGACCACATTG |
| 5634 | Va-PomAL183A-rv | CAATGTGGTCAAGgcTGCAACGGCC |
| 5635 | Va-PomAL183M-fwd | GGCCGTTGCAaTgTTGACCACATTG |

| | | |
|---|---|---|
| 5636 | Va-PomAL183M-rv | CAATGTGGTCAAcAtTGCAACGGCC |
| 5637 | Va-PomAL183F-fwd | GGCCGTTGCAtTtTTGACCACATTGTATG |
| 5638 | Va-PomAL183F-rv | CATACAATGTGGTCAAaAaTGCAACGGCC |
| 5639 | Va-PomAL184V-fwd | GGCCGTTGCACTCgTGACCACATTG |
| 5640 | Va-PomAL184V-rv | CAATGTGGTCAcGAGTGCAACGGCC |
| 5641 | Va-PomAT186A-fwd | GCACTCTTGACCgCATTGTATGGCG |
| 5642 | Va-PomAT186A-rv | CGCCATACAATGcGGTCAAGAGTGC |
| 5643 | Va-PomAA190I-fwd | CACATTGTATGGCattATCCTGTCC |
| 5644 | Va-PomAA190I-rv | GGACAGGATaatGCCATACAATGTG |
| 5645 | Va-PomAM195G-fwd | GATCCTGTCCAATggGGTGTTTTTCCC |
| 5646 | Va-PomAM195G-rv | GGGAAAAACACCccATTGGACAGGATC |
| 5647 | Va-PomAV196F-fwd | CCAATATGtTcTTTTTCCCTATTGCGG |
| 5648 | Va-PomAV196F-rv | CCGCAATAGGGAAAAAgAaCATATTGG |
| 5649 | Va-PomAR232A-fwd | GCCAAAACCCGgcAGTGATCGATAG |
| 5650 | Va-PomAR232A-rv | CTATCGATCACTgcCGGGTTTTGGC |
| 5651 | Va-PotBG20V-fwd | CCGTTATGGATGGtGACATTCGCAG |
| 5652 | Va-PotBG20V-rv | CTGCGAATGTCaCCATCCATAACGG |
| 5653 | Va-PotBG20W-fwd | CCGTTATGGATGtGGACATTCGCAG |
| 5654 | Va-PotBG20W-rv | CTGCGAATGTCCaCATCCATAACGG |
| 5655 | Va-PotBT21A-fwd | GGATGGGGgCATTCGCAGATTTGATG |
| 5656 | Va-PotBT21A-rv | CATCAAATCTGCGAATGcCCCCATCC |
| 5657 | Va-PotBF22Y-fwd | GATGGGGACATaCGCAGATTTGATG |
| 5658 | Va-PotBF22Y-rv | CATCAAATCTGCGtATGTCCCCATC |
| 5659 | Va-PotBD24A-fwd | GATGGGGACATTCGCAGcTTTGATGTC |
| 5660 | Va-PotBD24A-rv | GACATCAAAgCTGCGAATGTCCCCATC |
| 5661 | Va-PotBL25F-fwd | GGACATTCGCAGATTTtATGTCGCTGC |
| 5662 | Va-PotBL25F-rv | GCAGCGACATaAAATCTGCGAATGTCC |
| 5663 | Va-PotBS27A-fwd | CGCAGATTTGATGgCGCTGCTGATGTG |
| 5664 | Va-PotBS27A-rv | CACATCAGCAGCGcCATCAAATCTGCG |
| 5665 | Va-PotBL28Q-fwd | GATTTGATGTCGCaGCTGATGTGTTTC |
| 5666 | Va-PotBL28Q-rv | GAAACACATCAGCtGCGACATCAAATC |
| 5667 | Va-PotBF32A-fwd | CTGCTGATGTGTgcCTTTGTTCTTCTG |
| 5668 | Va-PotBF32A-rv | CAGAAGAACAAAGgcACACATCAGCAG |
| 5669 | Va-PotBL35V-fwd | GTGTTTCTTTGTTgTTCTGCTCTCG |
| 5670 | Va-PotBL35V-rv | CGAGAGCAGAAcAACAAAGAAACAC |
| 5671 | Va-PotBL36A-fwd | CTTTGTTCTTgcGCTCTCGTTTTCTG |
| 5672 | Va-PotBL36A-rv | CAGAAAACGAGAGCgcAAGAACAAAG |
| 5673 | Va-PotBL36Q-fwd | CTTTGTTCTTCaGCTCTCGTTTTCTG |
| 5674 | Va-PotBL36Q-rv | CAGAAAACGAGAGCtGAAGAACAAAG |
| 5756 | Va-PomAS25C-fwd_rth | tGCATCGGCATGTTTGTCGATG |
| 5757 | Va-PomAS25C-rv_rth | GCCACCAAGCACCATCGCC |
| 5758 | Va-PomAR88A-fwd:rth | gccAAAGGTGGTTTTCTTGCTC |
| 5759 | Va-PomAR88A-rv_rth | CGCCGCATCGGCCATTTCC |
| 5760 | Va-PomAK89A-fwd_rth | gcgGGTGGTTTTCTTGCTCTTG |
| 5761 | Va-PomAK89A-rv_rth | ACGCGCCGCATCGGCCATTTC |
| 5762 | Va-PomAE96A-fwd_rth | cgGAGATGGAAATAAACAACAC |

| | | |
|---|---|---|
| 5763 | Va-PomAE96A-rv_rth | CAAGAGCAAGAAAACCACC |
| 5764 | Va-PomAM155A-fwd | gcGATTGGCACCTTGGTTGGTC |
| 5765 | Va-PomAM155A-rv | TCCCATCGCAGGAGCAACG |
| 5766 | Va-PotBF22Y-fwd_rth | aCGCAGATTTGATGTCGCTGC |
| 5767 | Va-PotBF22Y-rv_rth | ATGTCCCCATCCATAACGG |
| 6299 | Va-PomA_K60A-fwd | GGTGCGACAgccATTGCTGGCAAAGCC |
| 6300 | Va-PomA_T185A -fwd | CGTTGCACTCTTGgCCACATTGTATGGC |
| 6301 | Va-PomA_L249A-fwd | CGTGCCgcTGAGATTGACGAGTAACTTG |
| 6302 | Va-PomA_D252A-fwd | CTTGAGATTGcCGAGTAACTTGGAGAG |
| 6303 | Va-PomA_DeltaC -fwd_rth | TAACTTGGAGAGTCGTGATG |
| 6304 | Va-PomA_K60A-rv | GGCTTTGCCAGCAATggcTGTCGCACC |
| 6305 | Va-PomA_T185A -rv | GCCATACAATGTGGcCAAGAGTGCAACG |
| 6306 | Va-PomA_L249A-rv | CAAGTTACTCGTCAATCTCAgcGGCACG |
| 6307 | Va-PomA_D252A-rv | CTCTCCAAGTTACTCGgCAATCTCAAG |
| 6308 | Va-PomA_DeltaC -rv_rth | TTCATTGAGGTAGTTCTTCAAG |
| 6299 | Va-PomA_K60A-fwd | GGTGCGACAgccATTGCTGGCAAAGCC |
| 6300 | Va-PomA_T185A -fwd | CGTTGCACTCTTGgCCACATTGTATGGC |
| 6301 | Va-PomA_L249A-fwd | CGTGCCgcTGAGATTGACGAGTAACTTG |
| 6302 | Va-PomA_D252A-fwd | CTTGAGATTGcCGAGTAACTTGGAGAG |
| 6303 | Va-PomA_DeltaC -fwd_rth | TAACTTGGAGAGTCGTGATG |
| 6304 | Va-PomA_K60A-rv | GGCTTTGCCAGCAATggcTGTCGCACC |
| 6305 | Va-PomA_T185A -rv | GCCATACAATGTGGcCAAGAGTGCAACG |
| 6306 | Va-PomA_L249A-rv | CAAGTTACTCGTCAATCTCAgcGGCACG |
| 6307 | Va-PomA_D252A-rv | CTCTCCAAGTTACTCGgCAATCTCAAG |
| 6308 | Va-PomA_DeltaC -rv_rth | TTCATTGAGGTAGTTCTTCAAG |
| 6540 | p43_fwd | ATGATTGGCACCTTGGTTGG |
| 6541 | p43_rev | ATAATGACGAATGCAAAACC |
| 6542 | gBlock_p43_fwd | ggttttgcattcgtcattatGGCGATGGTGCTTGGTGG |
| 6543 | gBlock_p43_rev | ccaaccaaggtgccaatcatTCCCATCGCAGGAGCAAC |
| 6544 | p47_fwd | TCGCAGATTTGATGTCGCTG |
| 6545 | p47_rev | GCCATACAATGTGGTCAAGA |
| 6546 | gBlock_p47_fwd | tcttgaccacattgtatggcGCGATCCTGTCCAATATG |
| 6547 | gBlock_p47_rev | cagcgacatcaaatctgcgaATGTCCCCATCCATAACG |
| Sequencing primers | | |
| 3117 | 5'_pBAD24_seq_fw | cgggaccaaagccatgacaa |
| 440 | 3'-pTrc-seq-rv | ggcaaattctgttttatcagac |
| 5497 | pomA_Va_check-fwd | GCCTTCATGTTTAAAGCGGA |
| 5498 | pomB_Va_check-fwd | GTGTTTCTTTGTTCTTCTGC |
| gBlocks | | |

| | | |
|---|---|---|
| 128 | p43_gBlock | GGCGATGGTGCTTGGTGGCAGCATCGGCATGTTTGTCGATGTCACGT CGATCCTTATTGTCGTTGGTGGCTCAATATTCGTCGTGTTGATGAAGT TCACAATGGGACAGTTTTTGGTGCGACAgccATTGCTGGCAAAGCCT TCATGTTTAAAGCGGATGAACCCGAAGACCTGATCGCAAAAATTGTG GAAATGGCCGATGCGGCGCGTAAAGGTGGTTTTCTTGCTCTTGAAGAG ATGGAAATAAACAACACATTCATGCAGAAAGGCATTGATCTACTGGTT GATGGCCATGATGCCGACGTTGTGAGAGCGGCACTCAAAAAAGACATC GCGCTTACGGATGAACGACATACGCAAGGTACTGGTGTATTTCGCGCCT TTGGCGACGTTGCTCCTGCGATGGGA |
| 129 | p47_gBlock | GCGATCCTGTCCAATATGGTGTTTTTCCCTATTGCGGATAAACTTTCTC TTCGCCGTGACCAAGAAACGCTAAATCGCCGTTTGATCATGGATGGCG TATTAGCGATTCAAGATGGCCAAAACCCGCGAGTGATCGATAGTTACT TGAAGAACTACCTCAATGAATAACTTGGAGAGTCGTGATGGATGATGA AGATAACAAATGCGATTGTCCGCCACCTGGCCTCCCGTTATGGATGGG GACAT |
